# Closed-loop auditory stimulation targeting alpha and theta oscillations during REM sleep induces phase-dependent power and frequency changes

**DOI:** 10.1101/2024.03.03.582907

**Authors:** Valeria Jaramillo, Henry Hebron, Sara Wong, Giuseppe Atzori, Ullrich Bartsch, Derk-Jan Dijk, Ines R. Violante

## Abstract

**Background:** Alpha and theta oscillations characterize the waking human electroencephalogram (EEG) and can be modulated by closed-loop auditory stimulation (CLAS). These oscillations also occur during rapid eye movement (REM) sleep, but whether they can be modulated by CLAS is not known.

**Objective:** Investigate whether CLAS can modulate alpha and theta oscillations during REM sleep in a targeted phase-dependent manner.

**Methods:** We recorded high-density EEG during an extended overnight sleep period in 18 healthy young adults. Auditory stimulation was delivered during both phasic and tonic REM sleep in alternating 6 s ON and 6 s OFF windows. During the ON windows, stimuli were phase-locked to four orthogonal phases of ongoing alpha or theta oscillations detected in a frontal electrode (Fz).

**Results:** During ON windows, the four orthogonal phases of ongoing alpha and theta oscillations were targeted with high accuracy. Alpha and theta CLAS induced phase-dependent changes in power and frequency at the target location. Frequency-specific effects were observed for alpha trough (speeding up) and rising (slowing down) and theta trough (speeding up) conditions. These phase-dependent changes of CLAS were observed during both REM sleep substages, even though the amplitude evoked by auditory stimuli which were not phase-locked was very much reduced in phasic compared to tonic REM sleep.

**Conclusions:** This study provides evidence that faster REM sleep rhythms can be modulated by CLAS in a phase-dependent manner. This offers a new approach to investigate how modulation of REM sleep oscillations affects the contribution of this vigilance state to brain function.

**Highlights:**

- REM sleep alpha and theta oscillations can be modulated using phase-locked CLAS
- Phase-dependent changes in power and frequency are observed in the target area
- Phase-dependent modulation occurs in phasic and tonic REM sleep

**Graphical Abstract:** 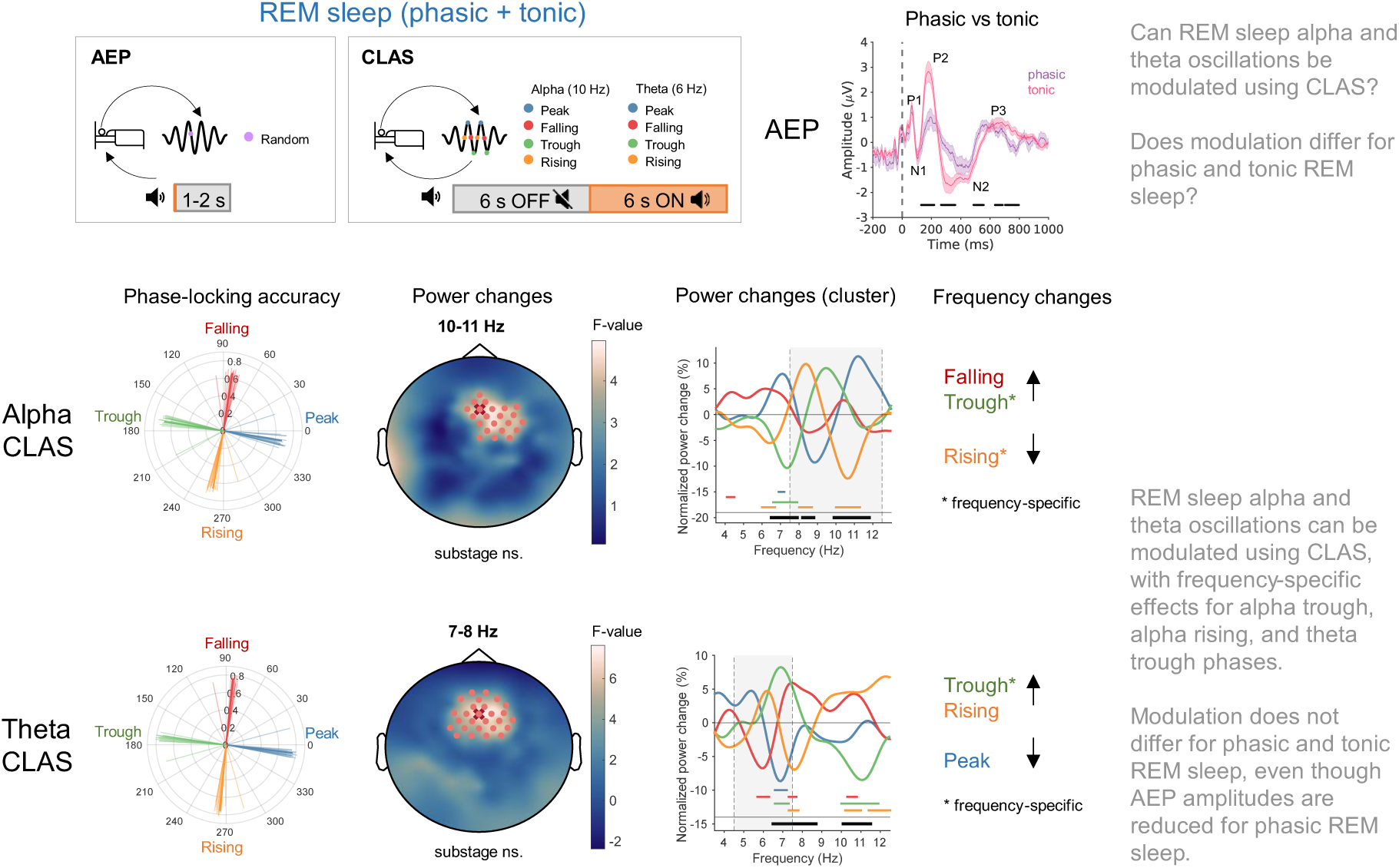

## Introduction

The brain produces a variety of oscillations which vary across vigilance states. During wakefulness, alpha (∼8-12 Hz) oscillations are dominant^1^ and thought to support various functions, including attention, perception, and memory^2,3^. The frequency of alpha oscillations varies and this is associated with brain function, with frequency reductions observed with cognitive decline in ageing and dementia^4–8^. In addition to alpha, theta (∼4-8 Hz) oscillations too can be observed during wakefulness and have been linked to working and episodic memory^9–11^.

Alpha and theta oscillations are likewise present during rapid eye movement (REM) sleep^12–18^. In fact, REM sleep is also termed paradoxical sleep because brain activity during this state is similar to wakefulness, but voluntary muscle activity is inhibited^19^. Alpha and theta oscillations are however more transient during REM sleep than wakefulness, and much less is known about their nature and function during REM sleep^12–18^.

REM sleep is further characterised by the occurrence of periods displaying rapid eye movements, muscle twitches, and increases in heart rate and breathing rate (phasic REM) which are interleaved with more quiescent periods (tonic REM)^20^. Phasic and tonic REM sleep appear to be very different brain states: Compared to tonic, in phasic REM sleep, alpha power and connectivity are reduced^21,22^, oscillation frequency is increased^23^, and the arousal threshold is increased^24,25^. Furthermore, the amplitude of auditory evoked potentials (AEPs) is much smaller during phasic REM sleep^26–29^. Finally, the proportion of these substages are altered in clinical populations e.g. depression^30^ and post-traumatic stress disorder^31^. These differences between phasic and tonic REM sleep have led to a renewed interest in their functional roles.

We recently demonstrated that alpha oscillations during wakefulness can be modulated using phase-locked closed-loop auditory stimulation (CLAS)^32^. Crucially, modulation of these oscillations was dependent on the phase that was targeted^32^. Here, we aimed to test if alpha and theta activity during REM sleep can also be modulated using CLAS and whether any effects are dependent on target phase or REM sleep substage.

## Material and methods

### Participants

Data were collected in 19 participants, one was excluded due to technical issues. The 18 participants included in the analysis were aged between 18-30 years (23.0 ± 2.0 [SD] y.o.; 7 males; 1 non-binary). Exclusion criteria are listed in **Suppl. Materials**. Participants gave written informed consent. The study conformed to the Declaration of Helsinki and received ethical approval from the University of Surrey’s Ethics Committee.

### Experimental protocol

Participants were instructed to keep a regular sleep-wake rhythm 7 days before the overnight recording. Adherence was controlled by wrist actimetry (material specifications listed in **Suppl. Materials**) and sleep diaries. Overnight recordings took place in a temperature-controlled, sound-attenuated, windowless sleep laboratory at the Surrey Sleep Research Centre (SSRC). On the day of the experiment, participants arrived at the SSRC 3 hours before their habitual sleep time. Biopac electrodes were applied to the participant’s scalp and connected to the custom-made endpoint-corrected Hilbert Transform (ecHT) device that was used for phase-locking^33^: A signal electrode was placed on Fz, reference and ground electrodes on the right mastoid. Gold electrodes were attached to the chin and next to the eyes and disposable electrodes were applied to the chest according to the American Academy of Sleep Medicine (AASM) guidelines^34^. Biopac and gold electrodes were attached using low impedance electrode cream. A high-density EEG cap was adjusted to Cz, and electrodes were filled with electrolyte gel. Impedances were below 50 kΩ at the start of the experiment. High-density EEG data were referenced to Cz. All EEG recordings were sampled at 500 Hz. In-ear headphones were secured in participant’s ears using medical tape. Half an hour before sleep, the Karolinska Sleepiness Scale (KSS) and a Visual Analogue Mood Scale (VAMS) were administered (**Figure 1A**). Then, wake EEG was recorded during which AEPs (see “AEPs” section) were elicited. This was followed by a 10-hour sleep opportunity starting at the habitual sleep time. We used a 10-hour extended sleep period to substantially increase time in REM sleep^35,36^. The sleep EEG was continuously monitored by a sleep expert and whenever participants entered REM sleep auditory stimulation was initiated. Stimulation was stopped when participants transitioned to NREM sleep or if they woke up. Stimulation started with an AEP block with the same protocol as during wakefulness. This was followed by an alpha or theta CLAS block (see “CLAS” section). The order of the CLAS blocks was counter-balanced across participants, as was the order of the different phases within each block. 10 hours after lights off, participants were woken up, after which a structured interview probing dream experiences (DQ) and an auditory stimulation questionnaire (AQ) were administered. Participants were then offered breakfast. Half an hour after wake-up, the KSS and VAMS were administered. Finally, another AEP block was run. KSS, VAMS, DQ, and AQ results are summarised in **Suppl. Materials**.

**Figure 1.**
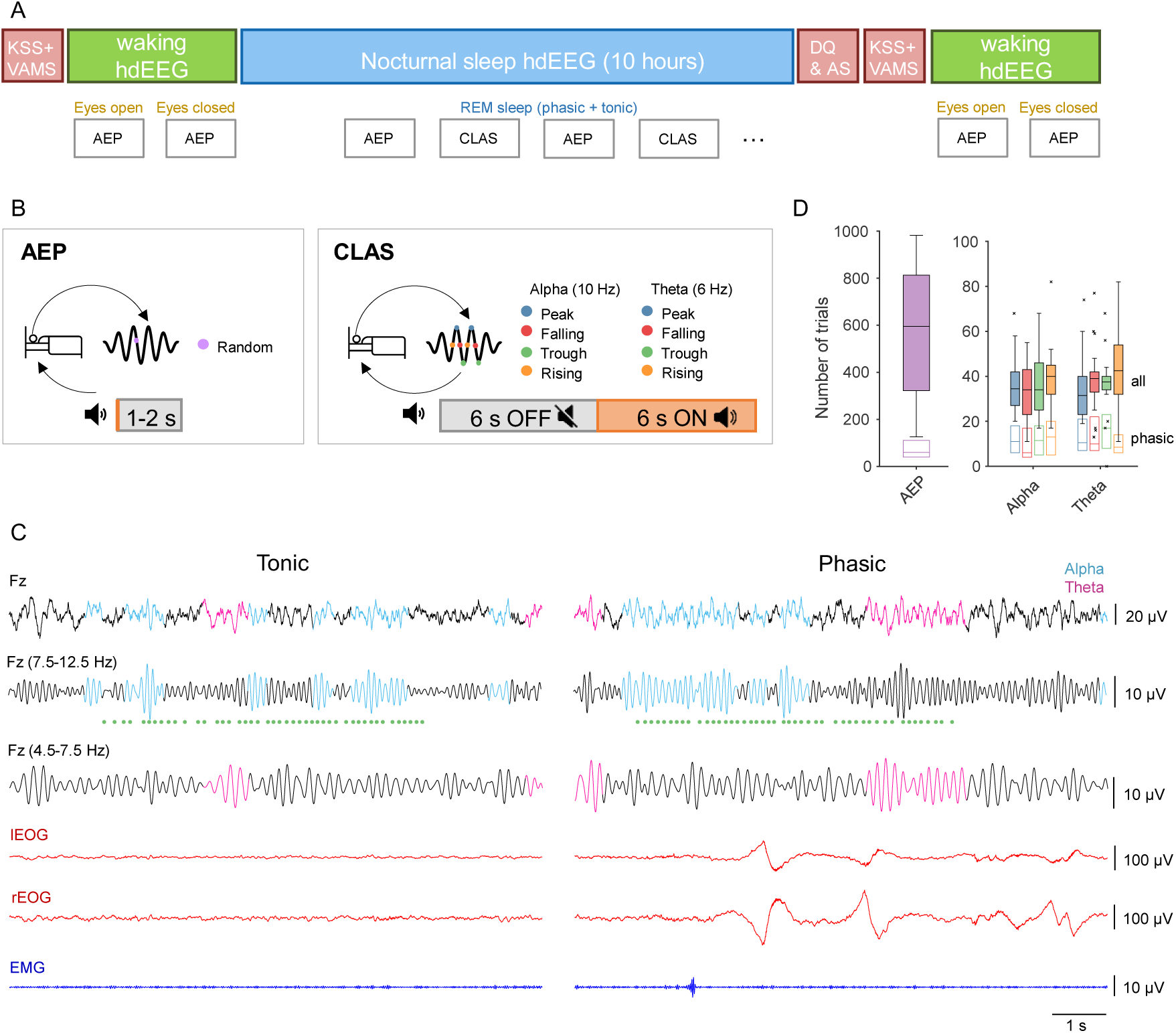
Experimental protocol. **A** In the evening, participants were fitted with a high-density EEG. After a questionnaire block comprising the KSS and VAMS, AEPs were recorded during wakefulness with eyes open and eyes closed. During the night, participants stayed in bed for 10 hours and AEPs and CLAS blocks were delivered during phasic and tonic REM sleep. In the morning, questionnaire blocks comprising DQ, AQ, KSS, VAMS were followed by AEP blocks during wakefulness with eyes open and eyes closed. **B** During AEP blocks, sounds were delivered every 1-2 seconds (not phase-locked) and at 3 different volumes. If a phasic 1-s segment occurred in the 6 s before or after the sound the trial was classified as phasic, otherwise it was classified as tonic. During CLAS blocks, sounds were delivered phase-locked to alpha or theta-filtered signal in 6 s ON (stimulation) and 6 s OFF (no stimulation) windows. 4 different phases were targeted and the order of stimulation was counter-balanced (for phase and frequency). If a phasic 1-s segment occurred in the 6 s before or after the start of the ON window the trial was classified as phasic, otherwise it was classified as tonic. **C** Example of tonic and phasic REM sleep segment in which CLAS was delivered. Showing EEG activity at channel Fz (phase-locking channel) for the broadband (top row) and filtered in alpha and theta frequency (second and third rows, respectively). Alpha and theta oscillations were detected post-hoc using eBOSC and are highlighted in colour. Green dots represent times at which auditory stimulation was delivered. Bottom rows show left (l) and right (r) EOG channels. Last row shows EMG activity. **D** Number of trials. Left panel shows number of AEP trials in total (filled purple bar) and for phasic REM sleep (bar with white fill and purple outline). Right panel shows the number of 12 s ON-OFF blocks for each CLAS condition in total (filled bar) and for phasic REM sleep (bar with white fill and coloured outline). To evaluate differences in the number of trials across phase conditions, a linear mixed-effects model with number of trials as dependent variable, condition as fixed factor and participant as random factor, was calculated. The condition factor was not significant (p > 0.05) indicating no differences in the number of trials for the different phase conditions. KSS = Karolinska Sleepiness Scale. VAMS = Visual Analogue Mood Scale. hdEEG = high-density EEG. DQ = Dream Questionnaire. AS = Auditory Stimulation Questionnaire. AEP = Auditory evoked potentials. eBOSC = extended Better Oscillation detection.

#### AEPs

AEPs were elicited during wakefulness and REM sleep by delivering single pulses of auditory stimuli (20 ms, pink noise) separated by 1-2 s of silence (1.5 ± 0.3 s) (**Figure 1B**). Stimuli were delivered at 50, 55, and 60 dB. During wakefulness (evening and morning), the protocol was run with eyes open (EO) and then repeated with eyes closed (EC). Auditory stimuli were not phase-locked to the ongoing brain oscillations.

#### CLAS

During CLAS blocks, the signal at the target electrode (Fz) was filtered in the alpha (7.5-12.5 Hz) or theta (4.5-7.5 Hz) range and the phase of the filtered signal was computed in real-time using the ecHT as in^32,33^. Stimuli (20 ms, pink noise) were delivered phase-locked to 4 different phases of the ongoing oscillation (peak, falling, trough, rising) in alternating 6 s ON and 6 s OFF windows (**Figure 1B**). The initial CLAS volume was set to the highest volume from which participants did not wake up during the first AEP block. If required, the volume was adapted (between 50-60 dB, was lowered if participants woke up and increased if they did not).

### EEG Preprocessing

EEG data were filtered, and sleep scoring performed as in^32^ and described in **Suppl. Materials**. REM sleep substages were scored in 1-s epochs (further information in **Suppl. Materials**). The signal of all channels was visually inspected and channels with poor quality were interpolated, and artefacts marked. An independent component analysis was run on the cleaned signal to identify and remove ocular and cardiac components. The signal was then re-referenced to the average across all electrodes.

### EEG Analyses

Power, oscillation detection, stimulation frequency, phase-locking accuracy, instantaneous frequency, and connectivity analyses were performed as in^32^ and described in **Suppl. Materials**.

### Statistics

Linear mixed-effects models were calculated for each electrode with statistical nonparametric mapping cluster correction to control for multiple comparisons^37^ (described in **Suppl. Materials**). Power, frequency, and connectivity changes were calculated as ratios between ON and OFF windows and log-transformed. To evaluate whether observed changes were significant t-tests were conducted. The significance level was set to 0.05. Values are presented as mean ± SD.

## Results

### Participants displayed phasic and tonic REM sleep during auditory stimulation

Participants slept on average 8.3 ± 0.8 hours out of which they spent 23.2 ± 4.4 % in REM sleep (**Table 1**). Participants displayed phasic and tonic REM sleep without signs of arousal during auditory stimulation (**Figure 1C**).

**Table 1.** Sleep parameters (N = 18).

|  | **Mean** | **SD** |
| --- | --- | --- |
| Total sleep time (TST, h) | 8.3 | 0.8 |
| Sleep latency (min) | 15.3 | 11.5 |
| WASO (min) | 93.7 | 44.8 |
| Sleep efficiency (%) | 82.3 | 7.8 |
| NREM (% TST) | 76.8 | 4.4 |
| REM (% TST) | 23.2 | 4.4 |
| Phasic REM (% REM) | 10.1 | 4.8 |
| Tonic REM (% REM) | 89.9 | 4.8 |

WASO = waking time after sleep onset. NREM = non-rapid eye movement sleep. REM = rapid eye movement sleep. SD = standard deviation.

Due to the natural lower proportion of phasic REM sleep fewer trials were obtained during this substage (**Figure 1D**). The number of trials did not differ between CLAS conditions (**Figure 1D; Suppl. Figure 1**). When comparing CLAS ON with OFF windows no difference in the proportion of phasic and tonic REM sleep was observed for most conditions – except when targeting the peak of alpha oscillations for which a slight increase in phasic REM sleep could be observed with stimulation (alpha peak: + 2.4 ± 3.4 %, p = 0.01; one-sample t-test).

### AEPs show major differences between phasic and tonic REM sleep

During wakefulness, power spectra showed significant differences between EC and EOs across a large range of frequencies, with the expected elevated alpha power during EC (**Figure 2A**). During REM sleep, the difference between substages was rather small with increased theta (∼3-5 Hz) and reduced high frequency (> 14 Hz) power for phasic compared to tonic REM sleep (**Figure 2B**). We next compared the responses to auditory stimulation for each brain state in the target area. Here, we observed a distinct pattern, such that there was a pronounced difference in the AEPs between phasic and tonic REM sleep but only minor differences between EO and EC (**Figure 2C-D**). Compared to phasic, tonic REM sleep showed greater AEP amplitude in three main time windows (P2: 132-218 ms; N2: 272-362 and 488-552 ms; and P3: 710-732 and 746-792 ms). The difference in AEPs remained after controlling for trial number between REM substages (**Suppl. Figure 2A**), circadian time and sleep pressure (**Suppl. Figure 2B**). We did not observe major differences in the AEPs in response to different sound volumes (**Suppl. Figure 2C**), therefore, this parameter was not considered in subsequent analyses.

**Figure 2.**
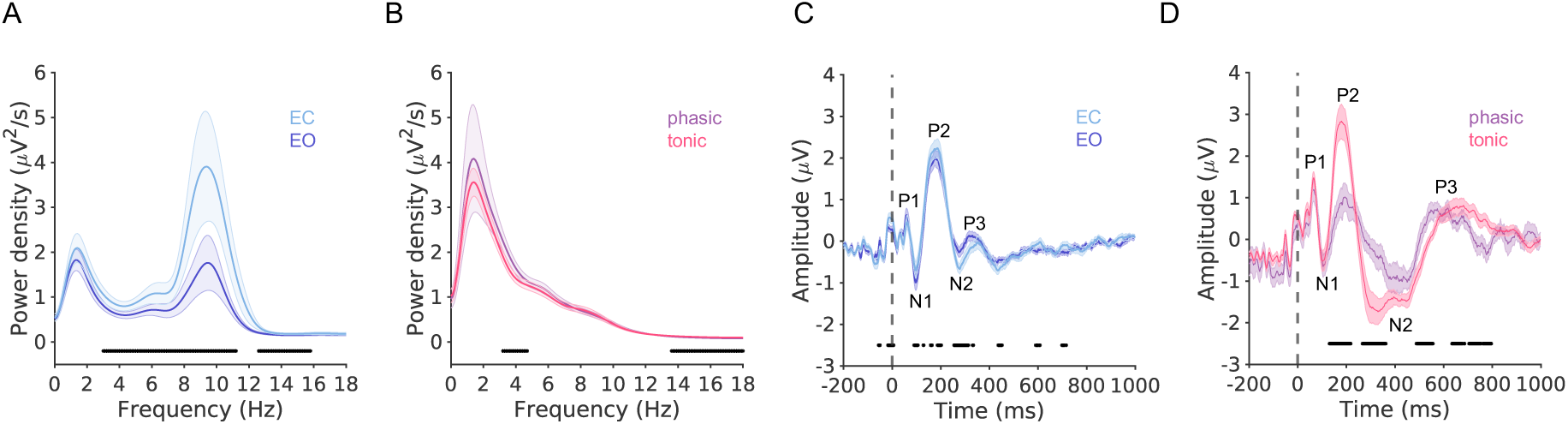
Power spectra and auditory evoked potentials (AEPs) differ for different vigilance states. **A** Power spectra for wakefulness (averaged for evening and morning) with eyes open and closed. **B** Power spectra for phasic and tonic REM sleep. **C** AEPs for wakefulness (averaged for evening and morning) during eyes open and closed. **D** AEPs for phasic and tonic REM sleep. Power spectra and AEPs were averaged across four electrodes closest to the phase-locking electrode. Vertical dashed lines denote stimulus onset. For each time point, a paired t-test was calculated to evaluate differences between the power spectra/AEPs. Significant time points are indicated with black bars at the bottom of the plot. EO = eyes open. EC = eyes closed.

### REM sleep alpha and theta oscillations can be targeted with high phase-locking accuracy

We first confirmed the presence of alpha and theta oscillations in all participants by visual inspection of the REM sleep EEG. Because these oscillations are transient, they are not easily identifiable in the power spectrum averaged across REM sleep epochs (**Figure 3A & 3F**, in target area, alpha peak identified in 6/18 and theta peak in 4/18 participants). Therefore, we employed the extended Better Oscillation detection (eBOSC) method^38^ to detect individual oscillations based on a power threshold that exceeds the 1/f background activity calculated over short time windows (30-s epochs). Using this method, we detected 1677 ± 633 alpha and 1315 ± 469 theta oscillations during REM sleep per participant at the target electrode (abundance, i.e., the duration of a rhythmic episode as a proportion of the epoch length, alpha: 0.19 ± 0.08, theta: 0.20 ± 0.05). Based on the distribution of detected oscillations, an alpha peak was identified in 13/18 and a theta peak in 12/18 participants.

**Figure 3.**
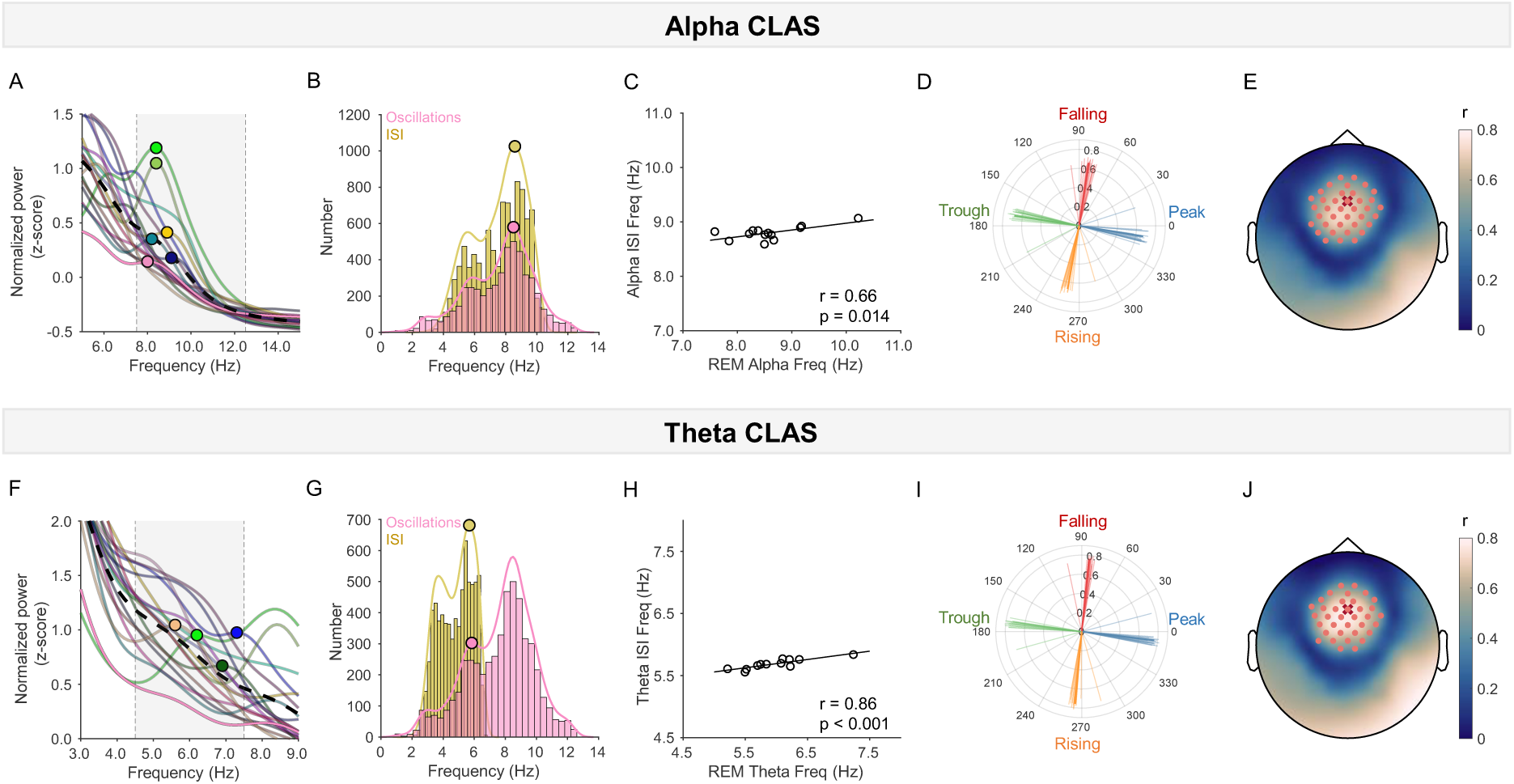
Validation of CLAS experiment. **A-E** refer to alpha CLAS whereas **F-J** refer to theta CLAS. **A and F:** Individual power spectra in REM sleep. Peaks are indicated with a dot. Note that not all participants have a peak in the alpha and/or theta range. **B and G:** Frequency distribution of oscillations detected during REM sleep (pink) and inter-stimulus-interval (ISI) frequency distribution (yellow) for one example participant. Peaks of the smoothed distributions (lines) are indicated with a dot. **C and H:** Association between REM oscillation frequency and frequency of stimulation across participants. Significance was assessed using Pearson correlations (only participants for which a peak in oscillation distribution could be detected were included, alpha: N=13, theta N=12). **D and I:** Circular plot showing the average phase targeted per participant (each line represents one participant). The length of the lines indicates the mean resultant vector length across all stimuli (ranges between 0 and 1 with higher values indicating higher phase-locking accuracy). The thick lines indicate the group average and resultant. **E and J:** Mean resultant vector length plotted for each electrode (averaged across all participants and phase conditions). For each phase condition and electrode, a V-test was computed across participants to evaluate if the data is non-uniformly distributed around the circle with a mean direction of the target phase (i.e. 0°, 90°, 180°, 270° for the different conditions). P-values were Bonferroni-corrected and only electrodes that were significant for all four phase conditions are indicated with pink dots. The electrode used for phase-locking is shown with a red cross.

To test if stimulation followed individual alpha and theta frequency, we determined individual peak frequency using the frequency distribution of the detected alpha and theta oscillations (**Figure 3B & 3G**). To estimate stimulation frequency, we calculated the interval between subsequent stimulations i.e. the inter-stimulus-interval (ISI), converted it to frequency and determined the peak in the ISI distribution. Stimulation frequencies were positively correlated with individual alpha and theta frequencies, yet the slopes were less than 1, indicating that stimulation only partially followed oscillations in these bands (**Figure 3C & 3H**). Stimuli were non-periodic (**Suppl. Figure 3**) and resultants for each condition, a measure of phase-locking accuracy, were high for the four targeted phases of alpha and theta oscillations with significant unimodal distributions (alpha: 0.67 ± 0.04, theta: 0.76 ± 0.03, all conditions z-stat > 11.8, p < 0.001) and average phase angles consistent across participants (alpha, phase angle deviation from target: -9.31° ± 7.13; theta: -5.79° ± 5.27), overall indicating high phase-locking accuracy (**Figure 3D & 3I**). Topographical analysis revealed highest phase-locking accuracy around the target electrode (**Figure 3E & 3J**). However, phase-locking was also comparatively high in the right mastoid area (where the reference electrode for the phase-locking was placed) and occipital areas. Since stimulation was delivered based on the signal filtered in the alpha or theta range, without online oscillation detection per se, we cannot exclude that part of the alpha phase-locked stimulation targeted theta oscillations, and vice versa (which could explain the imperfect frequency correlations). We therefore assessed phase-locking accuracy for the respective neighbouring band (**Suppl. Figure 4**). While resultants were largely reduced in the neighbouring band, the four targeted phases remained orthogonal to each other, but with opposite patterns compared to the targeted band, e.g. alpha peak stimulation resulted in theta trough stimulation.

### Alpha CLAS leads to phase-dependent modulation of alpha power and frequency

Having established that we can target alpha/theta oscillations in REM sleep in a phase-locked manner, we evaluated if phase-locked stimulation modulated EEG power, and if so whether this modulation was phase-dependent. We calculated power changes by comparing power in ON and OFF windows. For alpha CLAS, we found a significant effect of target-phase condition on power changes in frontal electrode clusters that were located around the target electrode (**Figure 4A**). This effect was restricted to 7-8 Hz and 10-11 Hz frequency bins (**Suppl. Figure 5**). This indicates that alpha power (and power at the border to the theta band) can be modulated by alpha phase-locked stimulation and that this effect depends on target phase. No significant effect of REM sleep substage (phasic vs tonic) and no interaction between substage and phase was found in these frequency bins, indicating that the modulation of power did not depend on the REM sleep substage (**Suppl. Figures 6-7**). However, substage, but not the interaction between substage and phase, was significant for slower (2-4 Hz) frequency bins, indicating that there was a phase-independent difference in power changes at slow frequencies for phasic and tonic REM sleep. Evaluating power changes at these slow frequencies for the two substages revealed a power reduction which was more pronounced for phasic REM sleep (**Suppl. Figure 8**).

**Figure 4.**
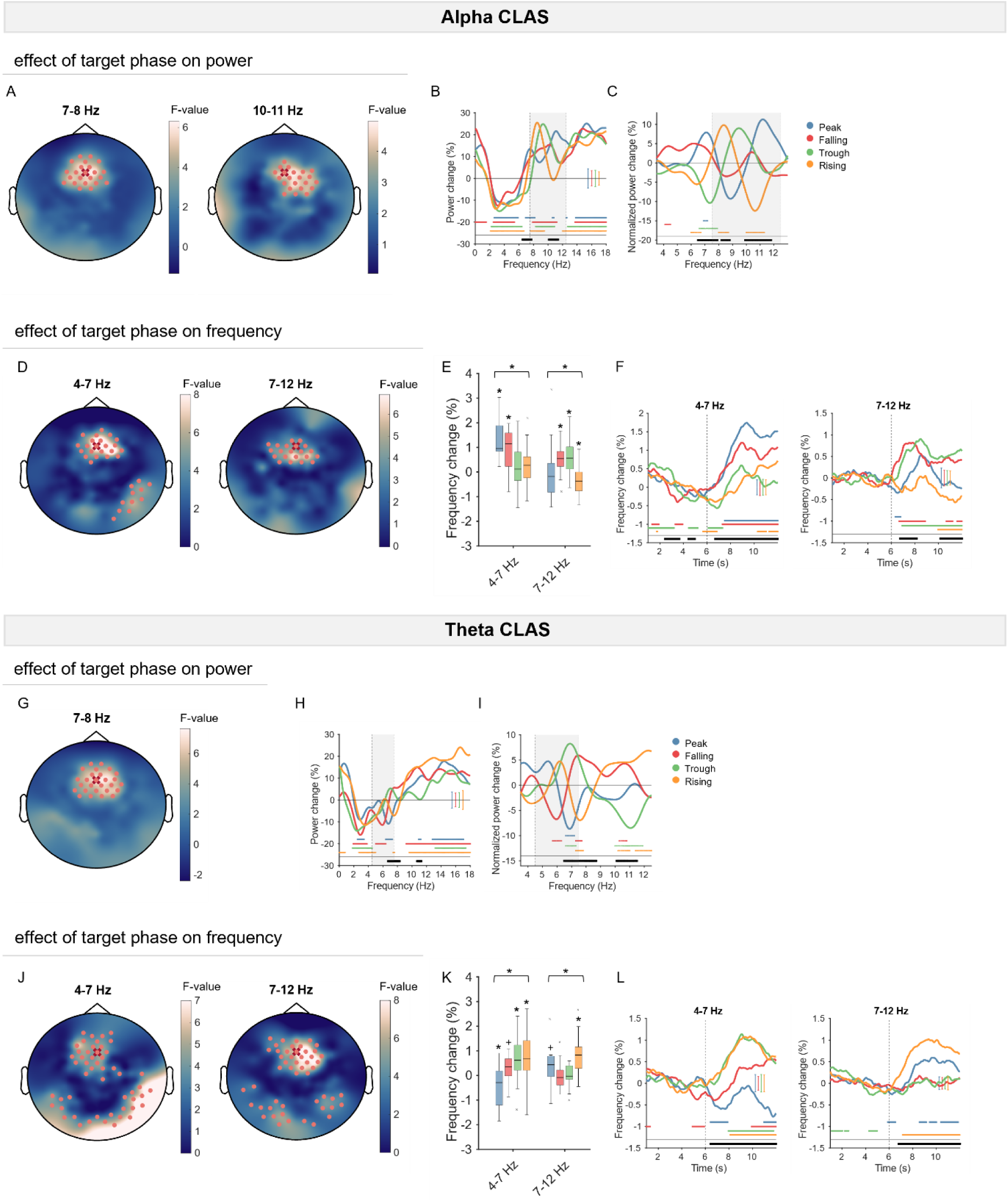
Phase-locked auditory stimulation induces power and frequency changes which depend on the target phase. **A-F** refer to alpha CLAS whereas **G-L** refer to theta CLAS. **A and G:** To evaluate differences in power changes across phase conditions, for each 1-Hz frequency bin between 1 and 16 Hz and for each electrode, linear mixed-effects models with power change as dependent variable, phase condition, substage, and the interaction between substage and phase condition as fixed factors, and participant as random factor were calculated. F-values for the fixed factor phase condition are plotted for each electrode. Significant electrodes are indicated with pink dots (cluster-corrected). The electrode used for phase-locking is shown with a red cross. Only frequency bins with significant electrode clusters are plotted here (see **Suppl. Figure 5 & 12** for all frequency bins). **B and H:** Power changes for each phase condition averaged across significant electrodes in **A and G**. Each line represents the average across participants for that phase condition, the error bar shows the averaged standard error of the mean across all frequency bins. To evaluate differences in power changes across phase conditions, linear mixed-effects models with power change as dependent variable, phase condition, substage, and the interaction between substage and phase condition as fixed factors, and participant as random factor were calculated. Significant frequency bins are indicated with black bars at the bottom of the plot. In addition, for each phase condition, significant changes were assessed using a one-sample t-test. Significant frequency bins are indicated with the colour corresponding to each phase condition at the bottom of the plot. Grey area indicates targeted frequency band i.e. frequency range in which the signal was filtered for phase-locking. **C and I:** Power changes in **B and H** were normalized with the average across all phase conditions. The same statistics as in **B and H** was performed using the normalized power values. **D and J:** To evaluate differences in frequency changes across phase conditions, linear mixed-effects models with frequency change as dependent variable, phase condition, substage, and the interaction between substage and phase condition as fixed factors, and participant as random factor were calculated. F-values for the fixed factor phase condition are plotted for each electrode. Significant electrodes are indicated with pink dots (cluster-corrected). **E and K:** Frequency changes for each phase condition averaged across significant electrodes in **D and J.** To evaluate differences in frequency changes across phase conditions, linear mixed-effects models with frequency change as dependent variable, phase condition, substage, and the interaction between substage and phase condition as fixed factors, and participant as random factor were calculated. A significant effect of phase condition is indicated with a black bar on top of the plot. Moreover, significant (p < 0.05) changes in a one-sample t-test are indicated with stars whereas trends (p < 0.1) are denoted with plus signs for each phase condition. **F and L:** Frequency changes for each time point during the ON window relative to the OFF window for each phase condition averaged across significant electrodes in **D and J**. Data was smoothed using a 2 s moving average (only including samples before the current sample). Each line represents the average across participants for that phase condition, the error bar shows the averaged standard error of the mean across all time points. The same statistics as in **B and H** was performed but for each time point (instead of frequency bin) and using frequency change (instead of power change) as dependent variable. Vertical dashed lines denote stimulation onset.

To better understand how the stimulation changes power across the spectrum, we assessed power changes averaged across the significant electrodes (**Figure 4B**). We found a general decrease in power for slower (2-7 Hz) and an increase for higher (> 12 Hz) frequencies across all phase conditions suggesting that the sound stimulation by itself i.e. without the phase-locking, induces phase-independent power changes. Importantly, the effect of phase condition remained significant only within the alpha band and at the border to the theta band indicating a phase-dependent modulation of alpha/theta power on top of the unspecific power changes. We further examined the phase-independent power changes across the scalp by averaging power across all phase conditions. This analysis revealed that these power changes were not only unspecific in terms of frequency but also location (**Suppl. Figure 9)**. To further examine the power changes that are phase-dependent, we eliminated the unspecific effects by normalizing to the averaged power changes across all phase conditions (**Figure 4C**). This analysis showed that power was increased at slightly different frequencies within the alpha band and decreased at neighbouring frequencies depending on the phase of stimulation. Such a pattern can often be observed when frequency changes occur. We therefore next assessed changes in oscillation frequency. It should be noted that power changes also extended into the neighbouring theta band, and thus in addition to alpha frequency changes we also evaluated changes in theta frequency.

To investigate changes in frequency we computed the instantaneous frequency within the alpha and the theta band for each electrode and compared ON and OFF stimulation windows. We observed a significant effect of phase condition on alpha and theta frequency changes in a frontal cluster around the target electrode (**Figure 4D**). In addition, a right occipital cluster was identified for the modulation of theta frequency. REM sleep substage and the interaction between condition and REM sleep substage were not significant (**Suppl. Figure 10**). These results indicate that alpha phase-locked stimulation modulated both alpha and theta frequency and that the modulation did not depend on the substage. We examined how the stimulation modulates frequency in more detail by averaging frequency changes across the significant electrodes. This analysis revealed a speeding up for the falling and trough conditions and a slowing down in alpha frequency for the rising condition (**Figure 4E**). In the theta frequency range, we detected a speeding up for peak and falling conditions. These frequency changes were sustained throughout the ON window for trough (alpha frequency), peak and falling conditions (theta frequency) and less sustained for the other conditions (**Figure 4F**).

Finally, we did not observe a prominent effect of alpha CLAS on connectivity, with significant effects of phase only present for phase-locking value (PLV) analysis, and only between the target region and the right mastoid (**Suppl. Figure 11**).

Taken together, these results demonstrate that REM sleep alpha power can be modulated using CLAS in a phase-dependent manner, and that this modulation is associated with changes in alpha frequency. In addition, phase-independent effects of stimulation as well as off-target effects on theta power and frequency were observed.

### Theta CLAS induces similar phase-dependent changes in theta power and frequency

We next evaluated changes in EEG power for theta CLAS by performing the same analyses as for alpha phase-locked stimulation. We found a significant electrode cluster in the 7-8 Hz frequency bin around the target electrode (**Figure 4G, Suppl. Figure 12**). Similar to alpha CLAS, no significant effect of substage or interaction between substage and condition was found for this frequency bin, but these factors were significant for other frequencies (**Suppl. Figures 13-14**). Again, this was due to larger power reductions at slower frequencies for phasic compared to tonic REM sleep but also larger increases at higher frequencies for tonic compared to phasic REM sleep (**Suppl. Figures 15-16**). Also, a frequency and location-unspecific decrease in power for slower frequencies (2-6 Hz) and an increase for higher frequencies (> 12 Hz) across all phase conditions was observed (**Figure 4H, Suppl. Figure 17**). When power changes were normalized with the averaged power changes across all phase conditions a similar pattern as for alpha CLAS emerged, but within the theta band: Power was increased at slightly different frequencies and decreased at neighbouring frequencies depending on the phase of stimulation (**Figure 4I**). In addition, power changes were also observed in the alpha band (∼10-12 Hz).

Topographical analysis of alpha and theta frequency changes revealed significant electrode clusters for the effect of phase condition for both alpha and theta frequency (**Figure 4J**). These were located around the target electrode, and in addition, in occipital electrode clusters. No significant effects of substage or the interaction between phase condition and substage were found (**Suppl. Figure 18**). Examining frequency changes across significant electrodes revealed a strikingly similar pattern to the one for alpha CLAS, but in the opposite direction: For theta frequency, a speeding up was observed for trough and rising conditions whereas a slowing down was detected for the peak condition (**Figure 4K**). For alpha frequency, a speeding up was found for the rising condition. Frequency changes were sustained throughout the ON window for trough (theta frequency) and rising (theta and alpha frequency) conditions and less sustained for the other conditions (**Figure 4L**). No significant effect of phase condition on PLV or PLI changes was found (**Suppl. Figure 19**).

To sum up, these results demonstrate that theta CLAS induces phase-dependent theta power and frequency changes. As for alpha CLAS, phase-independent effects of stimulation as well as off-target effects on alpha power and frequency were present.

## Discussion

In this study, we demonstrated that it is possible to perform alpha and theta phase-locked CLAS during REM sleep and that the stimulation induced phase-dependent changes in power and frequency. Although in an AEP paradigm we confirmed differences in the responsiveness to sounds for phasic and tonic REM sleep, the phase-dependent modulation of alpha and theta oscillations did not vary across these substages.

We first replicated previous studies showing pronounced differences in the amplitude of AEPs between wakefulness, phasic and tonic REM sleep^26–29^. Importantly, the difference in AEP amplitudes for phasic and tonic cannot be attributed to differences in circadian time or sleep pressure, since comparably small differences between AEPs were observed during wakefulness in the evening and morning. This difference in substage REM AEPs is especially striking considering the small differences in the power spectrum. Yet it confirms that phasic and tonic REM sleep represent distinctive states^20^.

We then demonstrated that alpha and theta oscillations during REM sleep can be modulated in a phase-dependent manner using CLAS. This modulation was most pronounced at the targeted frequency and location.

Previous studies have shown phase-dependent effects of stimulation targeting alpha or theta oscillations during wakefulness^32,39–46^ and slow waves during NREM sleep^47,48^. Some of these studies have reported phase-dependent effects on the perception of visual stimuli^39,40^ and phosphenes^41,42^ whereas others have found differences in motor evoked potentials^43–45^ and oscillatory power^46–48^. In our recent study, we showed that phase-locked CLAS targeting alpha oscillations during wakefulness does not only affect power but also frequency^32^. Here, we demonstrated remarkably similar changes in power and frequency when targeting REM sleep oscillations. Specifically, alpha frequency increased when targeting the trough and falling phase and decreased when targeting the rising phase of alpha oscillations. Theta frequency increased when targeting the trough and rising phase and decreased when targeting the peak of theta oscillations. Yet, some of these conditions also affected the neighbouring frequency band. To limit these, alpha CLAS trough (speed up) and rising (slow down) phases are preferable, whereas for theta CLAS the trough (speed up) is most promising to induce frequency-specific effects.

Surprisingly, considering the differences in the AEPs, these phase-dependent changes did not depend on REM sleep substage. Overall, the similar phase-dependent responses to auditory stimulation during wakefulness and REM sleep suggests that the oscillations occurring in these different states may share similar underlying network properties and thus respond similarly to external perturbation. We have previously proposed that CLAS-induced frequency changes may occur due to a phase-reset mechanism^32^. Such a mechanism is conceivable here; however, it remains an open debate if and how phase-reset can be differentiated from an evoked potential^49–53^.

While in our previous study during wakefulness we showed phase-dependent effects of stimulation on connectivity, here we report less definite results. Although we also found a phase-dependent effect of alpha CLAS on the PLV this was not confirmed using the PLI, a connectivity measure less likely to be confounded by volume conduction, but also more prone to disregard true phase relationships between different oscillators. Previous studies have shown increased connectivity in the alpha band during REM sleep^54,55^, and thus, it may be easier to modulate connectivity within a band that shows naturally higher synchronization across the brain.

A major difference to our study during wakefulness is that here we detected phase-independent reductions in low-frequency and increases in high-frequency power when performing phase-locked CLAS. Furthermore, these phase-independent changes differed for phasic and tonic REM sleep. This is consistent with the differences in the responsiveness to sounds per se that were also evident in the AEPs between these different states. Similar reductions in low frequency and increases in high frequency power in response to sound stimulation during REM sleep have been reported^56,57^. Interestingly, lower low frequency power and higher high frequency power have been associated with increased likelihood of conscious dream experiences and lucidity during REM sleep^58,59^ and sounds have been used to induce lucid dreaming^60^. Thus, sound stimulation during REM sleep may bring the brain into a more conscious state.

Our study design and analyses had limitations. First, no power thresholds for alpha or theta CLAS were used. Even though the phase-locking accuracy was low in the neighbouring frequency band, phase-targeting was consistent across participants and opposite to the targeted frequency band. As power and frequency tended to change in opposite directions for the neighbouring compared to the targeted band, we possibly simultaneously targeted both frequency bands. Alternatively, stimulation of one oscillation could have affected the other via an indirect phase-coupling mechanism. Further studies with more frequency-specific stimulation and computational modelling could disentangle these potential mechanisms. A second limitation is that the right mastoid was used as reference during phase-locking resulting in high phase-lock resultants in the right mastoid and occipital areas. Indeed, phase-dependent effects on frequency were also observed in these areas. Using a Laplacian reference during CLAS could avoid this issue in future studies. Thirdly, since our primary goal was to determine if alpha or theta oscillations during REM sleep can be modulated using CLAS, different stimulation conditions were delivered within the same night. Therefore, we were not able to assess if the stimulation affected sleep structure or functional outcome parameters. Yet, there are some indications for a functional role of REM sleep alpha/theta oscillations in dreaming^16–18,61^, learning and memory^57,62–68^, and cognitive function in ageing and dementia^17,61,69–73^. Future studies using fixed frequency-phase parameters per night can evaluate if stimulation affects such measures. Finally, CLAS was delivered throughout REM sleep, and although phasic and tonic stages were differentiated post-hoc, substage-specific stimulation may have identified differences in response to CLAS. In light that the reductions in low frequency power were more pronounced for phasic REM sleep, future studies should evaluate if these phase-independent effects can be attenuated by performing CLAS during tonic REM sleep only.

To conclude, our study demonstrates feasibility of modulating alpha and theta oscillations during REM sleep, such that power and frequency can be increased or decreased depending on which phase is targeted. This renders CLAS a promising tool to investigate relationships between these oscillations and functional outcome measures. The non-invasiveness and affordability of this tool allows to conduct larger-scale studies over prolonged periods of time in the future. Considering that alpha frequency becomes slower with ageing and dementia such a tool shows great therapeutic promise.

## Code and Data Availability

Code is available on https://gitlab.surrey.ac.uk/nemo/RSN. Raw data, sleep scoring, and data to make the figures will be made available on Zenodo (10.5281/zenodo.10663994) upon publication. Other processed data will be shared upon request.

## Author Contributions

VJ, HH, DJD, IRV: Conceptualization; VJ: Data curation; VJ: Formal analysis; VJ, DJD, IRV: Funding acquisition; VJ, SW, GA: Investigation; VJ: Project administration; UB, DJD, IRV: Resources; VJ, HH: Software; VJ, DJD, IRV: Supervision; VJ, HH: Visualization; VJ, DJD, IRV: Writing - original draft; VJ, HH, SW, UB, DJD, IRV: Writing - review & editing.

## Acknowledgements

VJ was supported by the Swiss National Science Foundation (P2EZP3_199918, P500PB_217827). IRV was supported by the Biotechnology and Biological Sciences Research Council (BB/S008314/1). HH was supported by the University of Surrey’s Doctoral College Scholarship Award. VJ, GA, DJD, UB were supported by UK the Dementia Research Institute (UKDRI-7005), Care Research and Technology Centre at Imperial College, London and the University of Surrey, Guildford. SW was supported by a UK DRI post-doctoral collaboration award to William Wisden, Nick Franks and DJD. UKDRI receives its funding from UK DRI Ltd, principally funded by the UK Medical Research Council, and additional funding partner Alzheimer’s Society. We would like to thank the members of the Surrey Sleep Research Centre, UKDRI CR&T Centre, Surrey Clinical Research Facility for their contribution to this work, in particular Katarina Jovic, Tegan Ward, and Jason Dai for their help in data collection. We thank the students at the Surrey School of Psychology Delia Lucarelli, Anika Thipaharan, Aliyah Khan, Radost Dimitrova, Chloe Moss, Elena Tomaselli, Mia Daniel, and Eliska Alderson, for their help in participant recruitment and data collection. We thank Kiran Ravindran for help in software development.

## Abbreviations

EEG: electroencephalogram
REM: rapid eye movement
CLAS: closed-loop auditory stimulation
AEP: auditory evoked potential
NREM: non-rapid eye movement
ecHT: endpoint-corrected Hilbert Transform
KSS: Karolinska Sleepiness Scale
VAMS: Visual Analogue Mood Scale
DQ: Dream Questionnaire
AQ: Auditory stimulation questionnaire
eBOSC: extended Better Oscillation detection
PLV: phase-locking value
PLI: phase lag index

