## Supplementary_Material for "Closed-loop auditory stimulation targeting alpha and theta oscillations during REM sleep induces phase-dependent power and frequency changes"

**Exclusion criteria**

Exclusion criteria were: Body mass index above 30, being pregnant, having acute or chronic disorders, taking medication which may have some cognitive effect, having severe scalp skin lesions, having undergone craniotomy or other acute neurosurgical intervention, having conducted shift work, maintained an irregular sleep/wake schedule or travelled through > 2 times zones during the preceding month, previously diagnosed sleep disorder, having a hearing impairment or hearing device.

**Material specifications**

| **Material** | **Manufacturer** |
| --- | --- |
| Actiwatch Spectrum Plus – wrist actimeter | Philips Respironics, US |
| 8 mm Biopac electrodes | Biopac, UK |
| Gold electrodes | CNSAC, Germany |
| Disposable ECG electrodes | Golmed, Germany |
| Low impedance electrode cream – used for Biopac and gold electrodes (Lic2) | CNSAC, Germany |
| High-density EEG cap (ActiCAP snap with 128 active electrodes) | Brain Products, Germany |
| Electrolyte gel (V20 electrolyte cream) – used for high-density electrodes | Brain Products, Germany |
| In-ear headphones (SE215) | Shure, US |
| Medical tape | Micropore, UK |

**Additional information on EEG preprocessing**

EEG data were preprocessed and analysed using Matlab 2020b (Mathworks, US). High-density EEG data were high-pass filtered (0.1 Hz followed by a 0.7 Hz filter) and notch filtered (50 Hz, 100 Hz) using the FieldTrip toolbox^1^. Sleep stages were scored in 30-s epochs according to standard AASM criteria^2^ using Domino (Somnomedics, Germany). For REM sleep epochs, phasic and tonic segments were scored in 1-s epochs based on the presence of a rapid eye movement in that segment. Electrodes with poor quality were interpolated using spheric linear interpolation.

**Additional information on EEG analyses**

Phasic and tonic percentages and trials

Phasic and tonic REM sleep percentages were computed based on 1-s segments. REM sleep AEP and CLAS trials were classified as phasic if there was at least one phasic 1-s segment in the 6 seconds preceding or following the stimulus (for AEP trials) or ON window onset (for CLAS trials). Trials that did not occur during REM sleep or contained artefacts were excluded.

Power analyses

For all power analyses, power was calculated in 1-s epochs (Hamming window, 0-40 Hz, in 0.1 Hz steps). using the *pwelch* function.

Oscillation detection

Oscillations were detected using the extended Better Oscillation detection (eBOSC) algorithm^3^. This method detects oscillations that exceed a power threshold that is determined based on a short time interval. Oscillation detection was carried out in 30-s epochs (1-30 Hz, in 0.25 Hz steps, 95 % power threshold). Only oscillations with at least 3 cycles and a duration of 300 ms or more were included in the analyses. For each participant, the number of waves and abundance with a frequency within the theta (4.5-7.5 Hz) and the alpha (7.5- 12.5 Hz) band was determined. The abundance represents the proportion of samples that are assigned to oscillatory activity within an epoch. For peak oscillation frequency estimation, the frequency distribution was smoothed using kernel density estimation (FWHM: 5.39 ± 1.94 Hz) before performing peak detection using the *findpeaks* function.

Stimulation frequency

Stimulation frequency was estimated by calculating the interval between subsequent stimuli (the inter-stimulus interval, ISI) and converting it to frequency (f = 1/ISI). For stimulation frequency estimation, the frequency distribution was smoothed using kernel density estimation (FWHM alpha: 5.17 ± 0.82 Hz, theta: 3.42 ± 0.50 Hz) before performing peak detection using the *findpeaks* function.

Phase-locking accuracy

Phase-locking accuracy was assessed using the high-density EEG data for each participant, electrode, and condition as follows: The phase at which each stimulus was delivered was estimated by running the ecHT algorithm offline on the last 1-s of EEG data before stimulation and extracting the phase at the end of the signal (i.e. at stimulus onset). The mean resultant vector length was then calculated across all stimuli. This gives a value between 0 and 1 with 0 indicating a uniform distribution around the circle and 1 a unimodal distribution i.e., perfect phase-locking.

Instantaneous frequency

Instantaneous frequency was estimated for each sample as proposed by Cohen^4^: The EEG signal was band-pass filtered in the alpha (7-12 Hz) or theta (4-7 Hz) range and Hilbert-transformed to obtain the phase angle timeseries. The difference between all sample points was taken to approximate the temporal derivate and smoothed using a median filter.

Connectivity

Connectivity was assessed using two different metrics: the phase-locking value (PLV) and the phase lag index (PLI) as proposed by Lachaux et al.^5^ and Stam et al.^6^ The PLI is less sensitive to volume conduction effects yet has also reduced sensitivity to detect true phase relationships between oscillators. To calculate these metrics the EEG signal was band-pass filtered for each channel in the delta (1-4 Hz), theta (4-7 Hz), alpha (7-12 Hz), and beta (13-30 Hz) bands and Hilbert-transformed. For each second, the phase-locking value (PLV) was calculated as the mean resultant vector length of all phase angles within that second. Similarly, the phase-lag index (PLI) was calculated for each second using the following formula: PLI = |mean(sign(Δphase))|. We kept our data in the sensor space because we do not have individual anatomical data (i.e., MRI scans) for each participant, hindering our ability to create realistic subject-specific head models. Furthermore, our closed-loop approach poses additional obstacles to source reconstruction, as it enforces a strong correlation between the auditory-evoked activity and the spontaneous activity. Nevertheless, a recent study comparing functional connectivity estimates from sensor and source spaces concluded that there is a strong correlation for the global connectivity between scalp- and source-level.^7^ In contrast, network topology (not investigated in our study) was only weakly correlated, albeit more accurate for functional connectivity metrics that limit the effects of volume conduction/signal leakage, such as PLI.

**Additional information on Statistics**

Circular statistics were performed using the CircStat toolbox^8^. Linear mixed-effects (lme) models were calculated using restricted maximum likelihood estimation of parameters using *fitlme* and analysis of variance tables of the fitted models were generated using *anova*. To correct for multiple comparisons in topographical analyses, the substage and the phase condition labels were shuffled randomly, and a lme was calculated for each electrode. This procedure was repeated 1000 times and for each permutation the maximal number of neighbouring electrodes with a p-value below 0.05 was determined to obtain a distribution of maximal cluster sizes. The cluster threshold was set to the 95th percentile of the maximal cluster size distribution.

**Questionnaire data**

When participants were asked if they could hear short bursts of sounds while they were sleeping or trying to fall asleep 17 out of 18 reported hearing the sounds. When asked about the last thing going through their mind prior to awakening, 3 participants reported hearing the sounds. While one of them had the impression that the sounds woke them up, one had thought-like experiences, and one had a vivid dream experience. Participants felt more energetic in the morning compared to the evening, but also felt less happy and less calm in the morning (**Suppl. Table 1**).

|  | evening | | morning | | evening vs morning | |
| --- | --- | --- | --- | --- | --- | --- |
|  | **Mean** | **SD** | **Mean** | **SD** | **t-value** | **p-value** |
| KSS | 6.9 | 0.2 | 3.5 | 1.3 | -11.30 | 2.5E-09 |
| VAMS energetic-sleepy | -50.6 | 11.0 | 33.9 | 33.3 | -10.40 | 9.0E-09 |
| VAMS happy-sad | 57.1 | 20.1 | 46.1 | 24.1 | 3.30 | 0.0042 |
| VAMS calm-tense | 61.9 | 22.8 | 45.8 | 28.0 | 2.44 | 0.0261 |

**Suppl. Table 1.** Overnight changes in subjective sleepiness and mood. The KSS ranges from 1-9, 9 is most sleepy; the VAMS ranges from -100 to 100, 100 is most energetic, happy, and calm.

KSS = Karolinska Sleepiness Scale, VAMS = Visual Analogue Mood Scale.

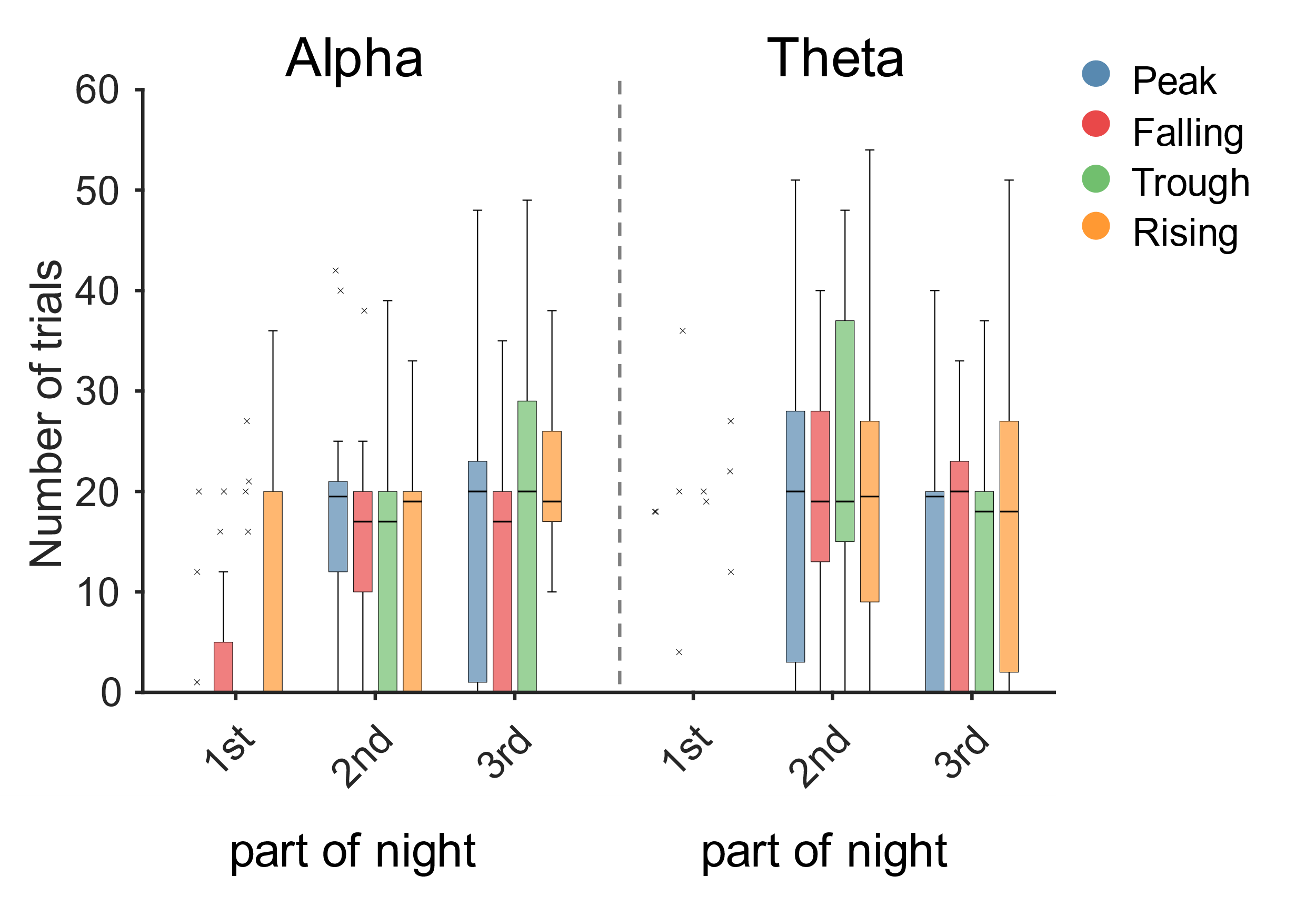

**Suppl. Figure 1.** Number of AEP trials for each phase condition and each third of the night. To evaluate differences in the number of trials across phase conditions, linear mixed-effects models with number of trials as dependent variable, condition as fixed factor and participant as random factor, were calculated. The condition factor was not significant (p > 0.05) in any of the models indicating no differences in the number of trials for the different phase conditions.

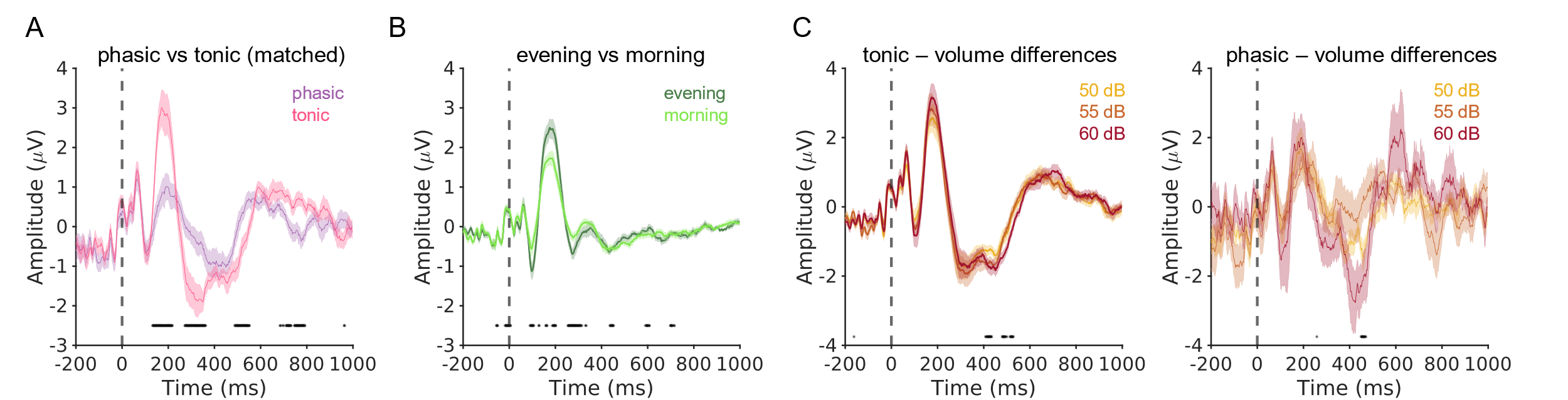

**Suppl. Figure 2.** **A** AEPs for phasic and tonic REM sleep when randomly selecting the same number of trials for tonic REM sleep as for phasic REM sleep. **B** AEPs for wakefulness in the evening and morning (averaged for eyes open and eyes closed). **A and B:** For each time point, a paired t-test was calculated to evaluate differences between the AEPs. **C** AEPs for different stimulation volumes for tonic (left) and phasic (right) REM sleep. Vertical dashed lines denote stimulus onset. For each time point, a linear mixed effects model with AEP amplitude as dependent variable, volume as fixed factor and participant as random factor was calculated. Significant time points are indicated with black bars at the bottom of the plot.

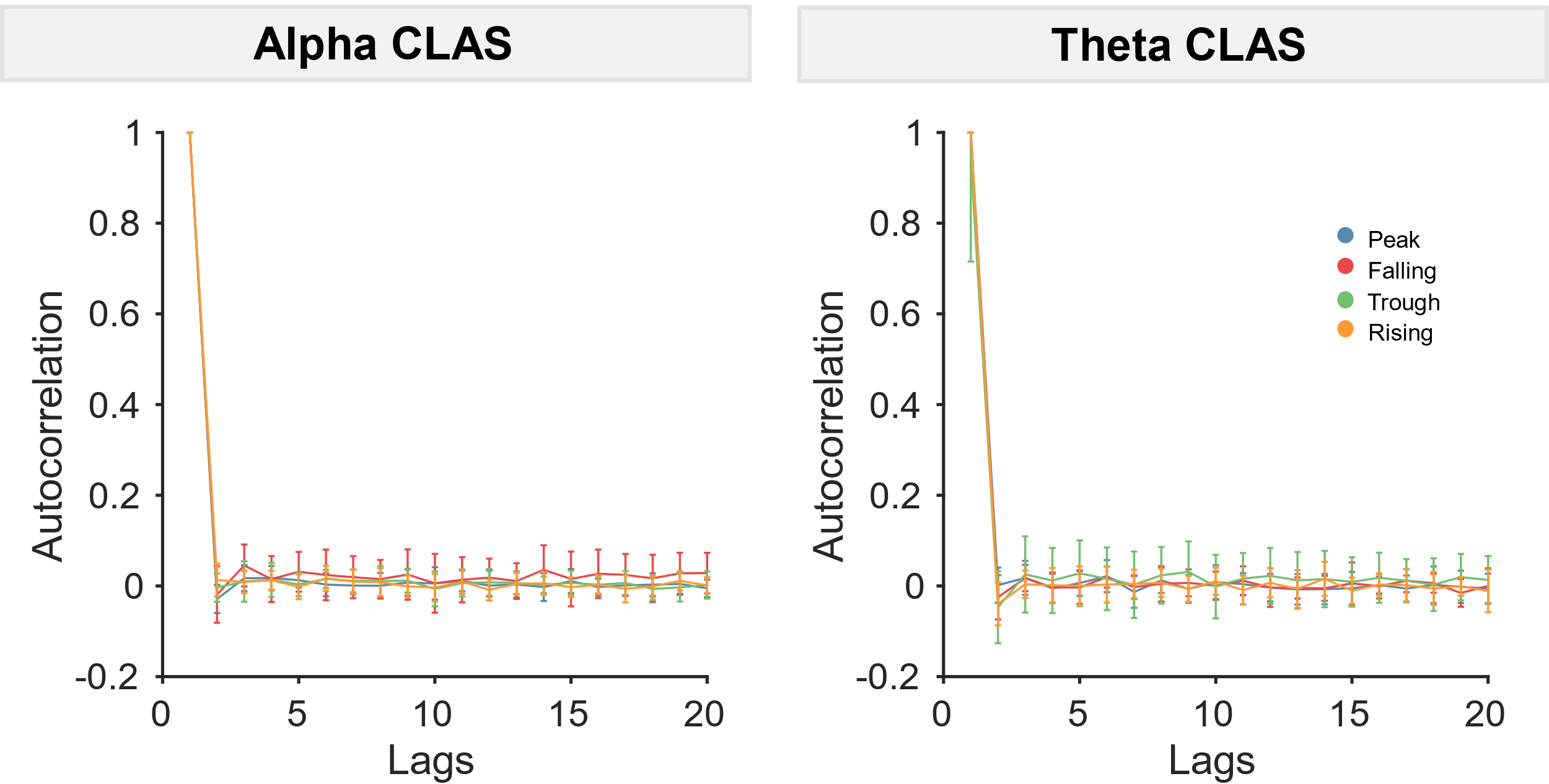

**Suppl. Figure 3.** Autocorrelation of ISIs. To evaluate periodicity of CLAS, ISIs were autocorrelated for each phase condition. Autocorrelation decreased rapidly after the second stimulus indicating that stimulation was non-periodic and therefore phase-dependent changes in frequency or power are unlikely to be explained by entrainment to a periodic stimulus.

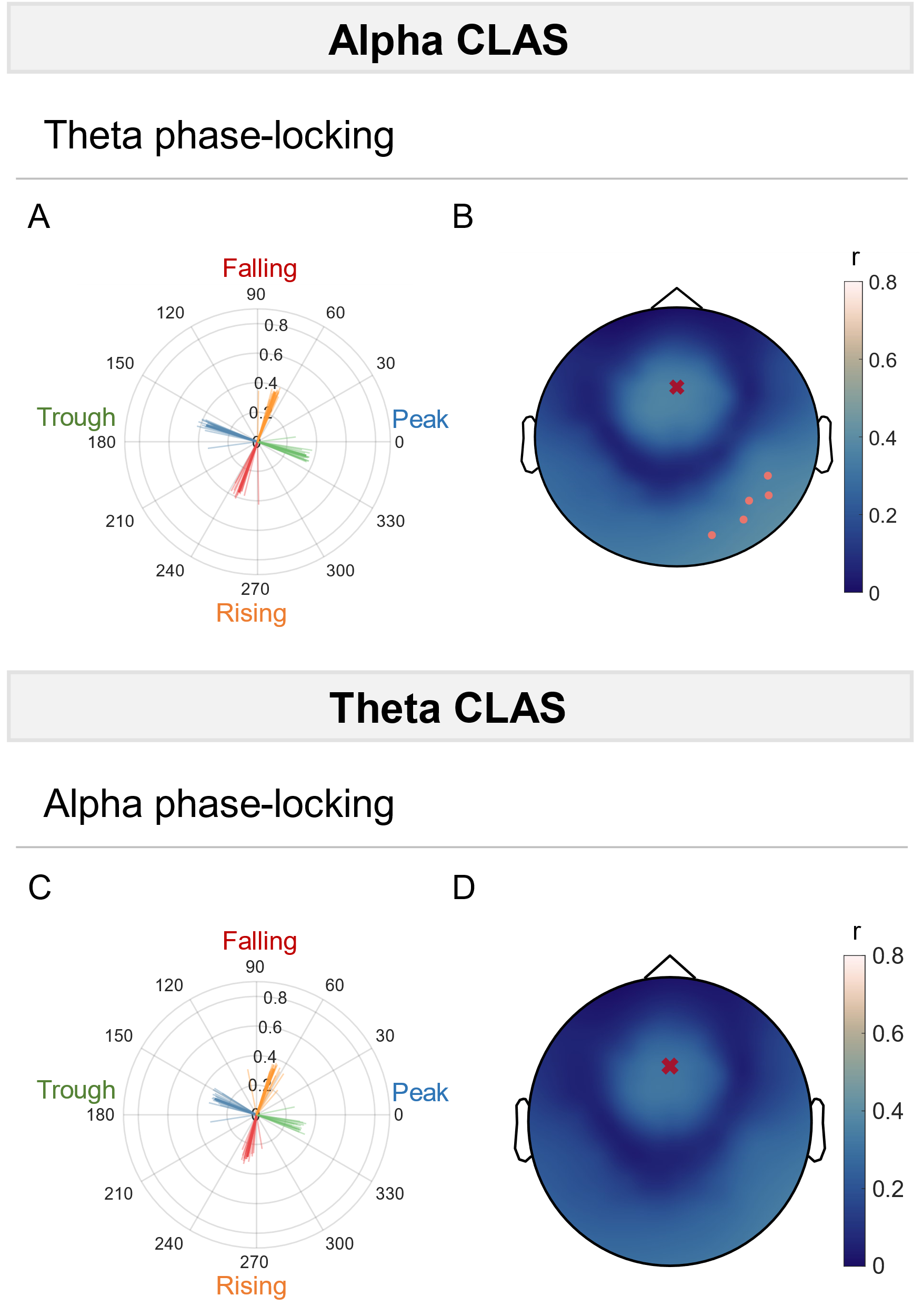

**Suppl. Figure 4. A and B:** Phase-locking accuracy for alpha CLAS when filtering the signal in the theta band at Fz in **A** and across all electrodes in **B**. **C and D:** Phase-locking accuracy for theta CLAS when filtering the signal in the alpha band at Fz in **C** and across all electrodes in **D**. For each phase condition and electrode, a V-test was computed across participants to evaluate if the data is non-uniformly distributed around the circle with a mean direction of the target phase (i.e. 0°, 90°, 180°, 270° for the different conditions). P-values were Bonferroni-corrected and only electrodes that were significant for all four phase conditions are indicated with pink dots. The electrode used for phase-locking is shown with a red cross.

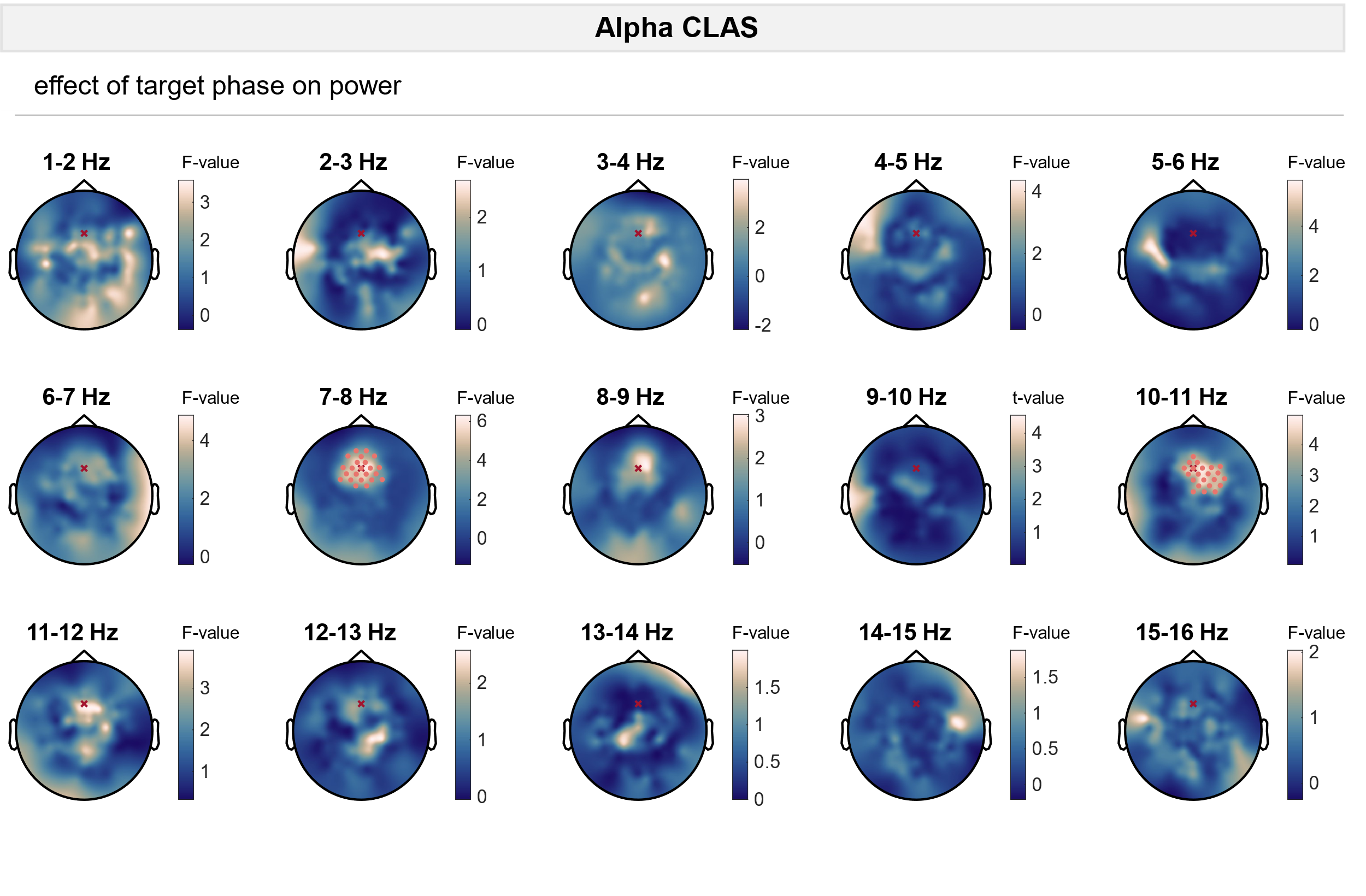

**Suppl. Figure 5.** To evaluate differences in power changes across phase conditions, for each 1-Hz frequency bin between 1 and 16 Hz and for each electrode, linear mixed-effects models with power change (ON vs OFF stimulation) as dependent variable, phase condition, substage, and the interaction between substage and phase condition as fixed factors, and participant as random factor were calculated. F-values for the fixed factor phase condition are plotted for each electrode. Significant electrodes are indicated with pink dots (cluster-corrected). The electrode used for phase-locking is shown with a red cross.

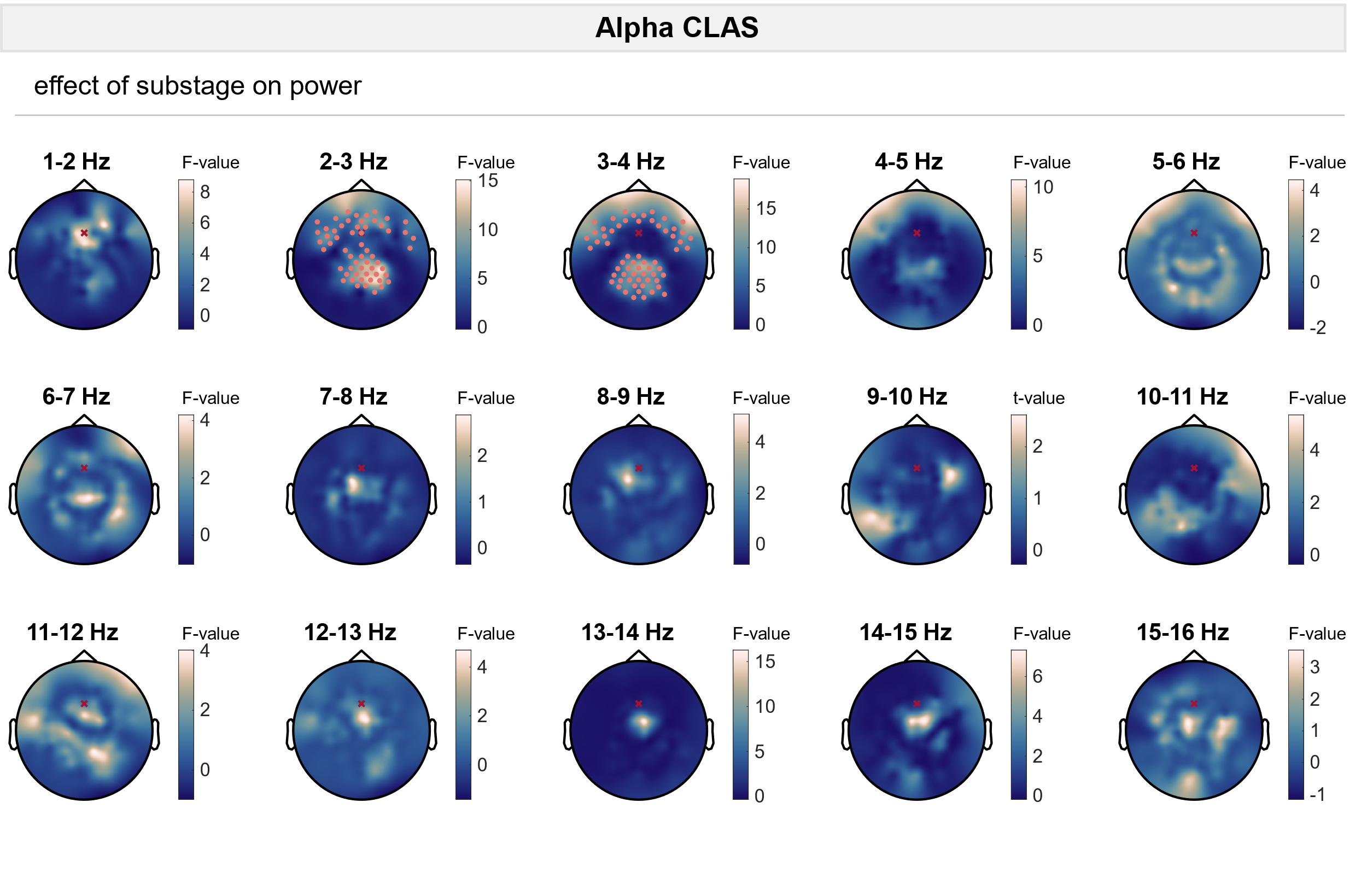
**Suppl. Figure 6.** To evaluate differences in power changes across phase conditions, for each 1-Hz frequency bin between 1 and 16 Hz and for each electrode, linear mixed-effects models with power change (ON vs OFF stimulation) as dependent variable, phase condition, substage, and the interaction between substage and phase condition as fixed factors, and participant as random factor were calculated. F-values for the fixed factor substage are plotted for each electrode. Significant electrodes are indicated with pink dots (cluster-corrected). The electrode used for phase-locking is shown with a red cross.

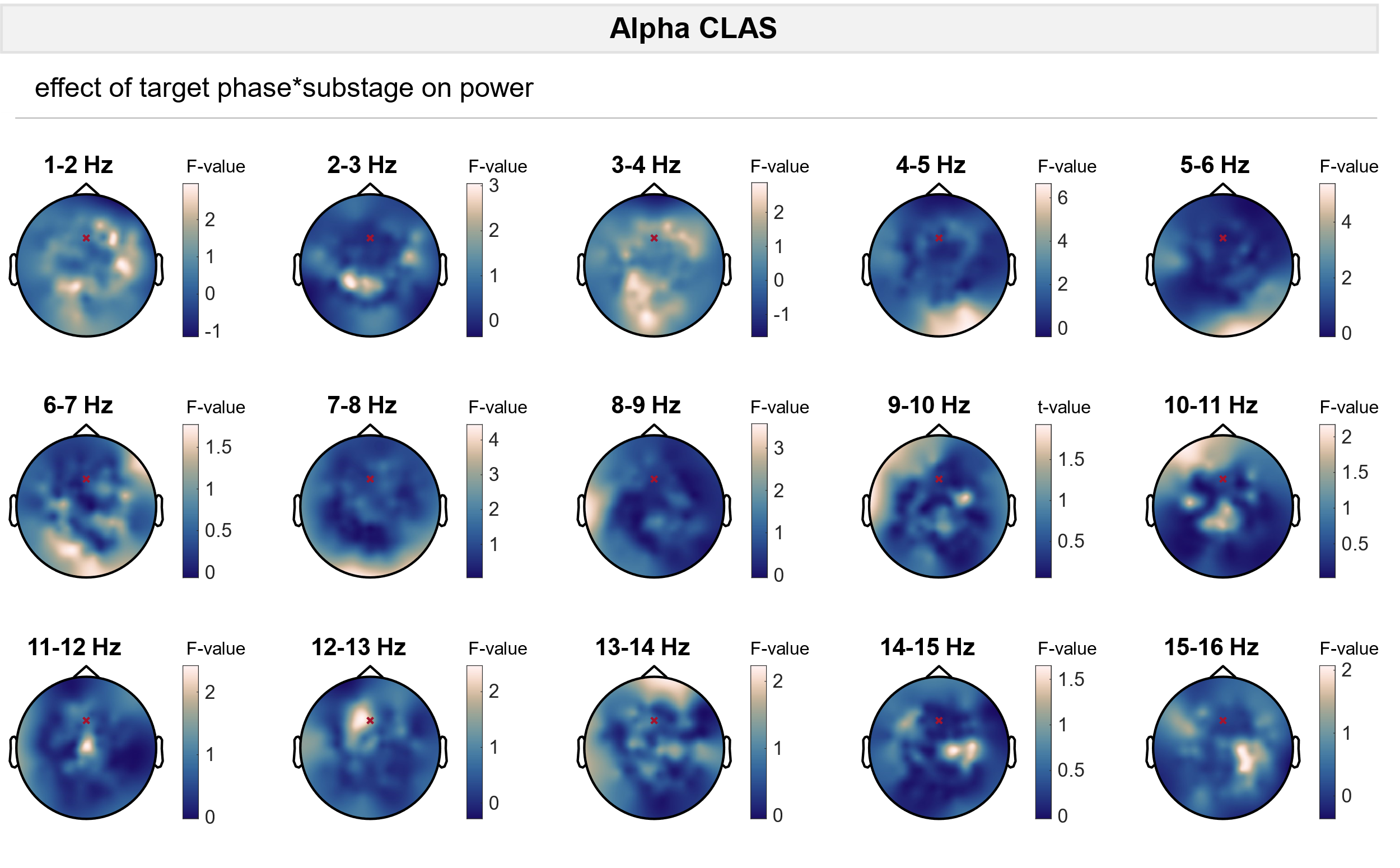

**Suppl. Figure 7.** To evaluate differences in power changes across phase conditions, for each 1-Hz frequency bin between 1 and 16 Hz and for each electrode, linear mixed-effects models with power change (ON vs OFF stimulation) as dependent variable, phase condition, substage, and the interaction between substage and phase condition as fixed factors, and participant as random factor were calculated. F-values for the interaction between phase condition and substage are plotted for each electrode. No significant (p < 0.05) electrode clusters were found. The electrode used for phase-locking is shown with a red cross.

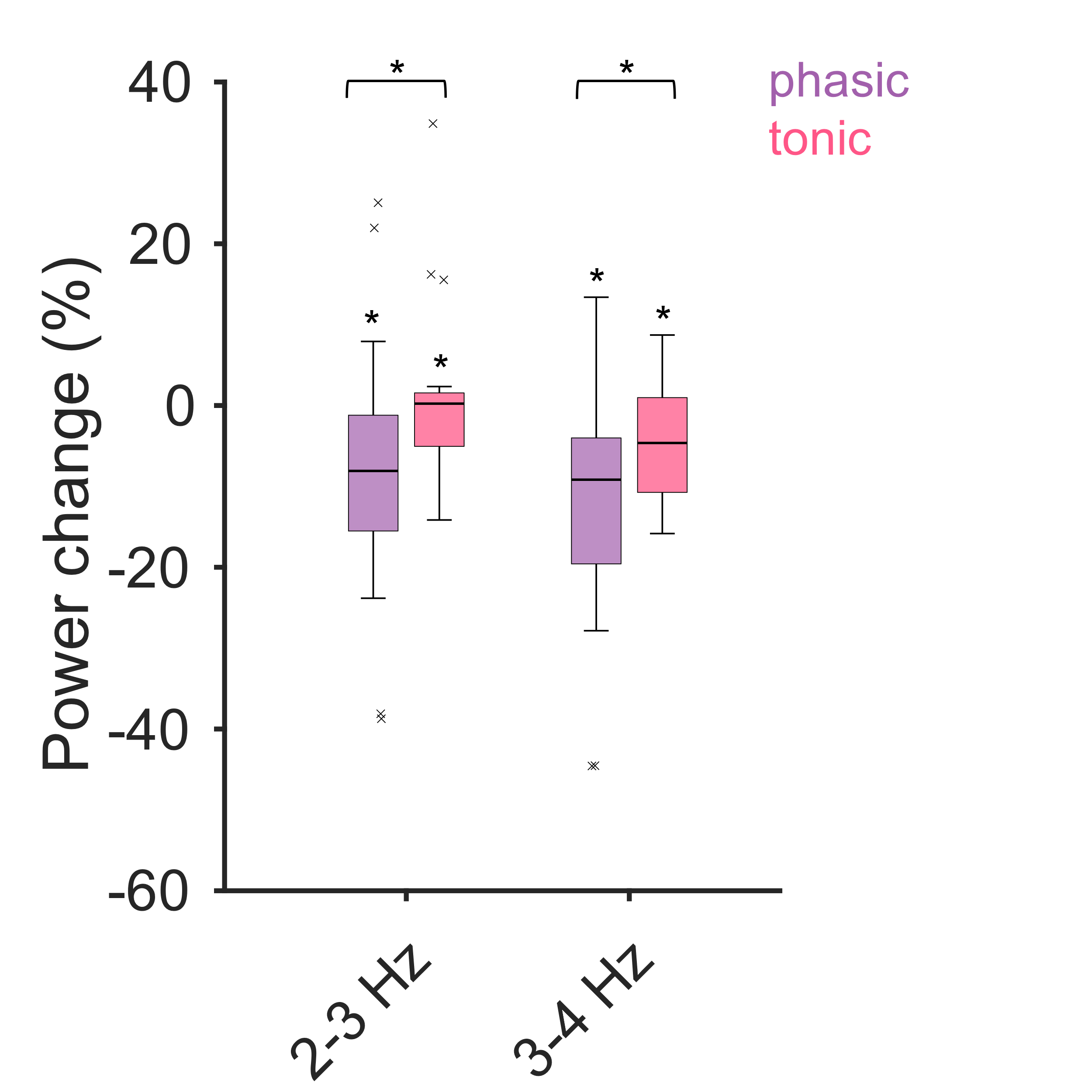

**Suppl. Figure 8.** Power changes for phasic and tonic REM sleep averaged across significant electrode clusters in **Suppl. Figure 6**. Significant differences between phasic and tonic REM sleep are indicated with black bars on top of the plot. Moreover, significant (p < 0.05) changes in a one-sample t-test are indicated with stars.

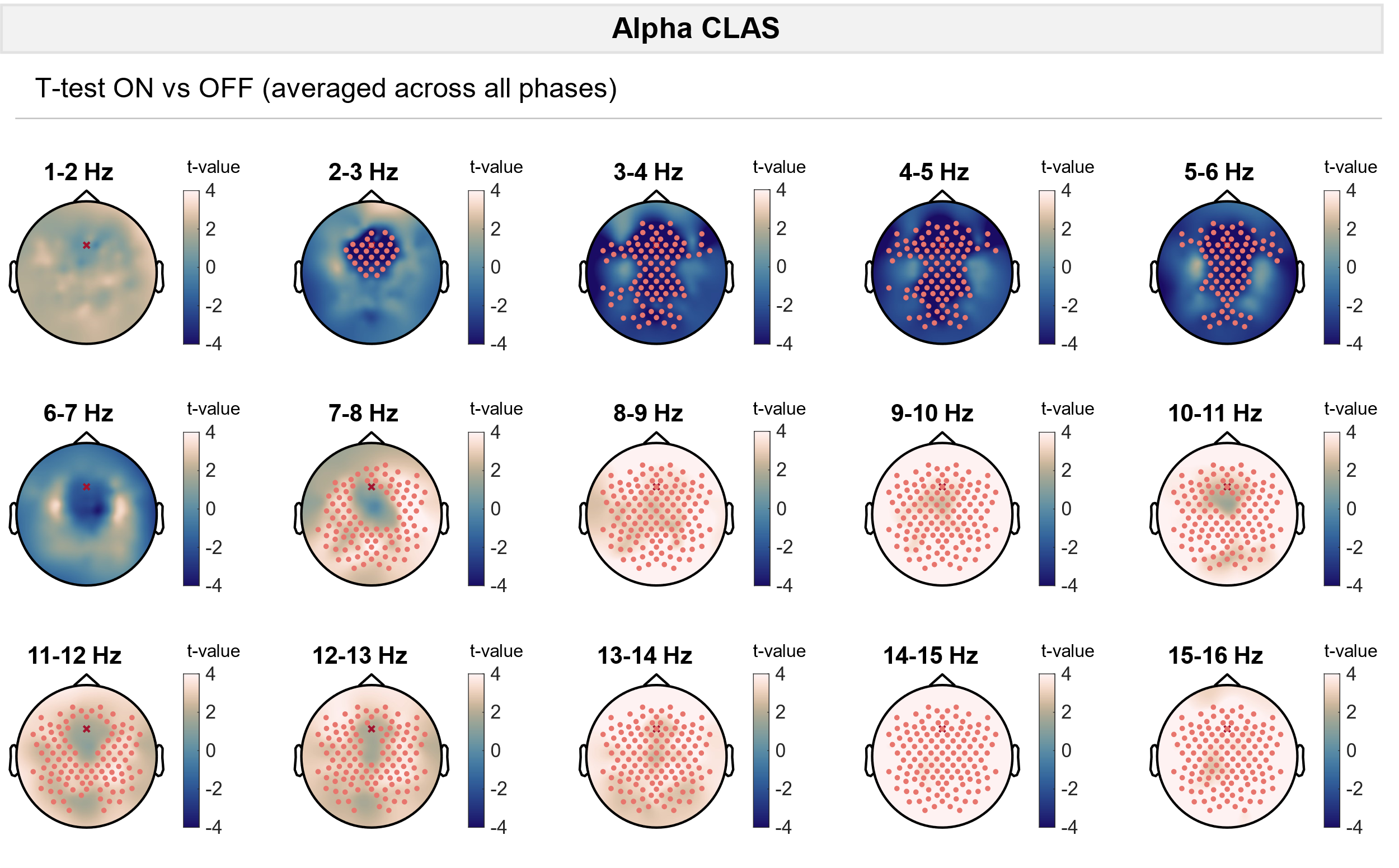
 **Suppl. Figure 9.** To evaluate phase-independent effects of stimulation, power changes were averaged across all phase conditions for ON and OFF windows. For each electrode, power in ON and OFF windows was compared using a paired t-test. Significant electrodes are indicated with pink dots (cluster-corrected). The electrode used for phase-locking is shown with a red cross.

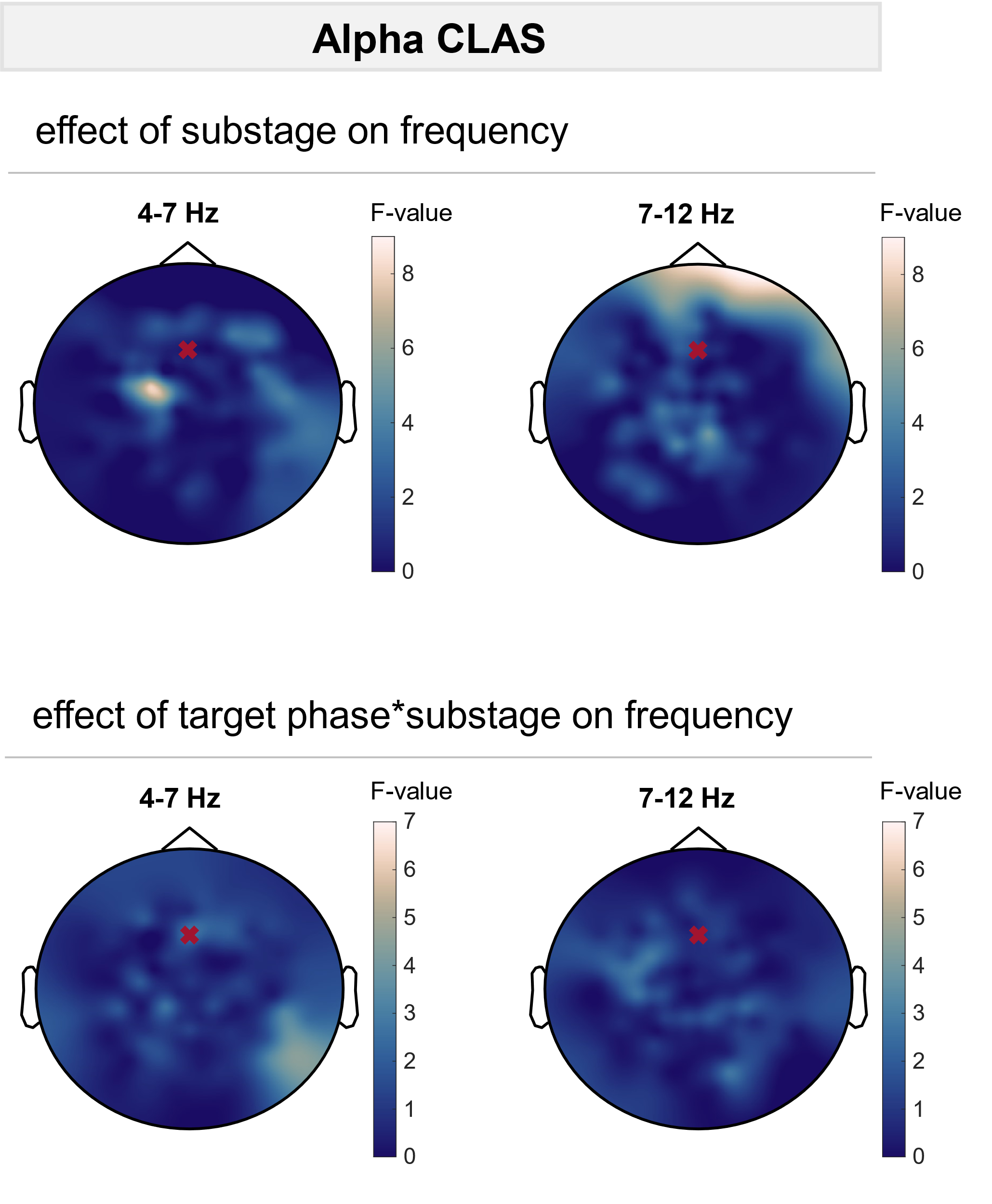

**Suppl. Figure 10.** To evaluate differences in frequency changes across phase conditions, linear mixed-effects models with frequency change as dependent variable, phase condition, substage, and the interaction between substage and phase condition as fixed factors, and participant as random factor were calculated. F-values for the fixed factor substage and the interaction between phase condition and substage are plotted for each electrode. No significant (p < 0.05) electrode clusters were found. The electrode used for phase-locking is shown with a red cross.

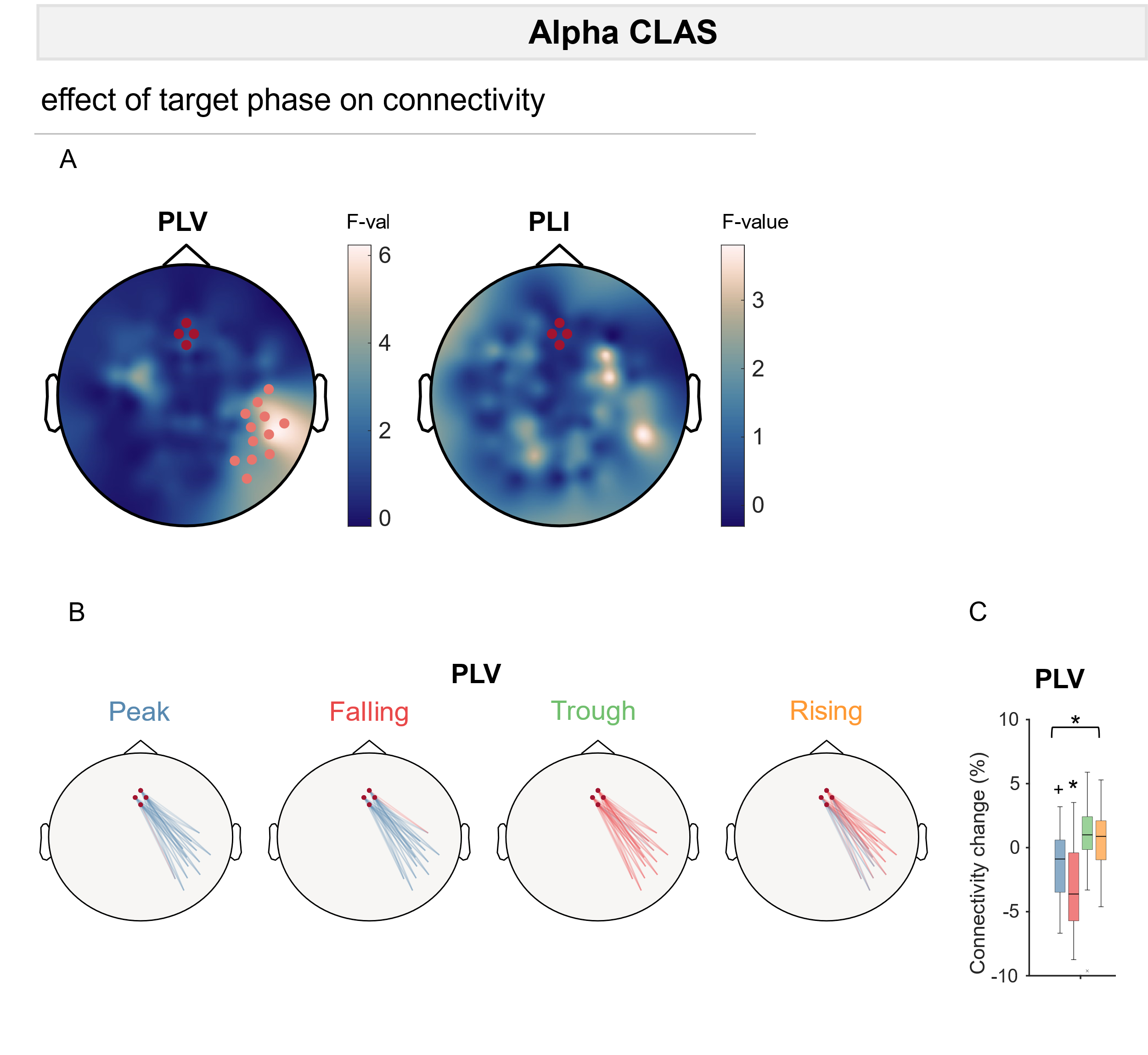

**Suppl. Figure 11.** Alpha CLAS may also induce phase-dependent changes in connectivity. Connectivity changes were assessed by calculating the PLV between the targeted region i.e. the seed region (red dots) and all other electrodes. **A** To evaluate differences in PLV and PLI changes across phase conditions, linear mixed-effects models with PLV or PLI change as dependent variable, phase condition, substage, and the interaction between substage and phase condition as fixed factors, and participant as random factor were calculated. F-values for the fixed factor phase condition are plotted for each electrode. Significant electrodes are indicated with pink dots (cluster-corrected). **B** PLV increases (red) and decreases (blue) between the seed region and the significant electrode cluster in **A** are indicated with lines. **C** PLV changes averaged across the significant electrode cluster in **A** for each phase condition. To evaluate differences in PLV changes across phase conditions, linear mixed-effects models with PLV change as dependent variable, phase condition, substage, and the interaction between substage and phase condition as fixed factors, and participant as random factor were calculated. A significant effect of phase condition is indicated with a black bar on top of the plot. In addition, significant (p < 0.05) changes in a one-sample t-test are indicated with stars.

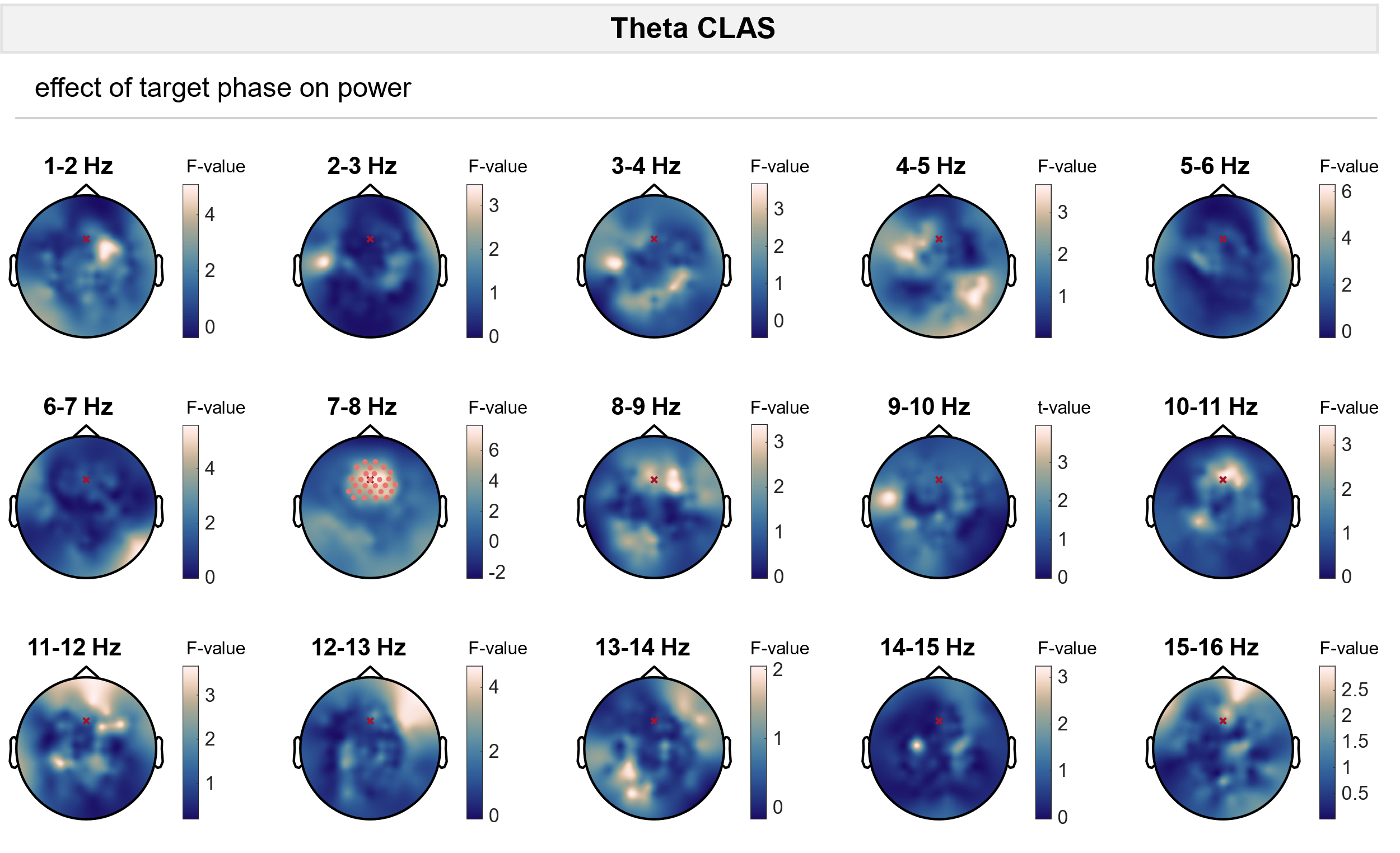

**Suppl. Figure 12.** To evaluate differences in power changes across phase conditions, for each 1-Hz frequency bin between 1 and 16 Hz and for each electrode, linear mixed-effects models with power change (ON vs OFF stimulation) as dependent variable, phase condition, substage, and the interaction between substage and phase condition as fixed factors, and participant as random factor were calculated. F-values for the fixed factor phase condition are plotted for each electrode. Significant electrodes are indicated with pink dots (cluster-corrected). The electrode used for phase-locking is shown with a red cross.

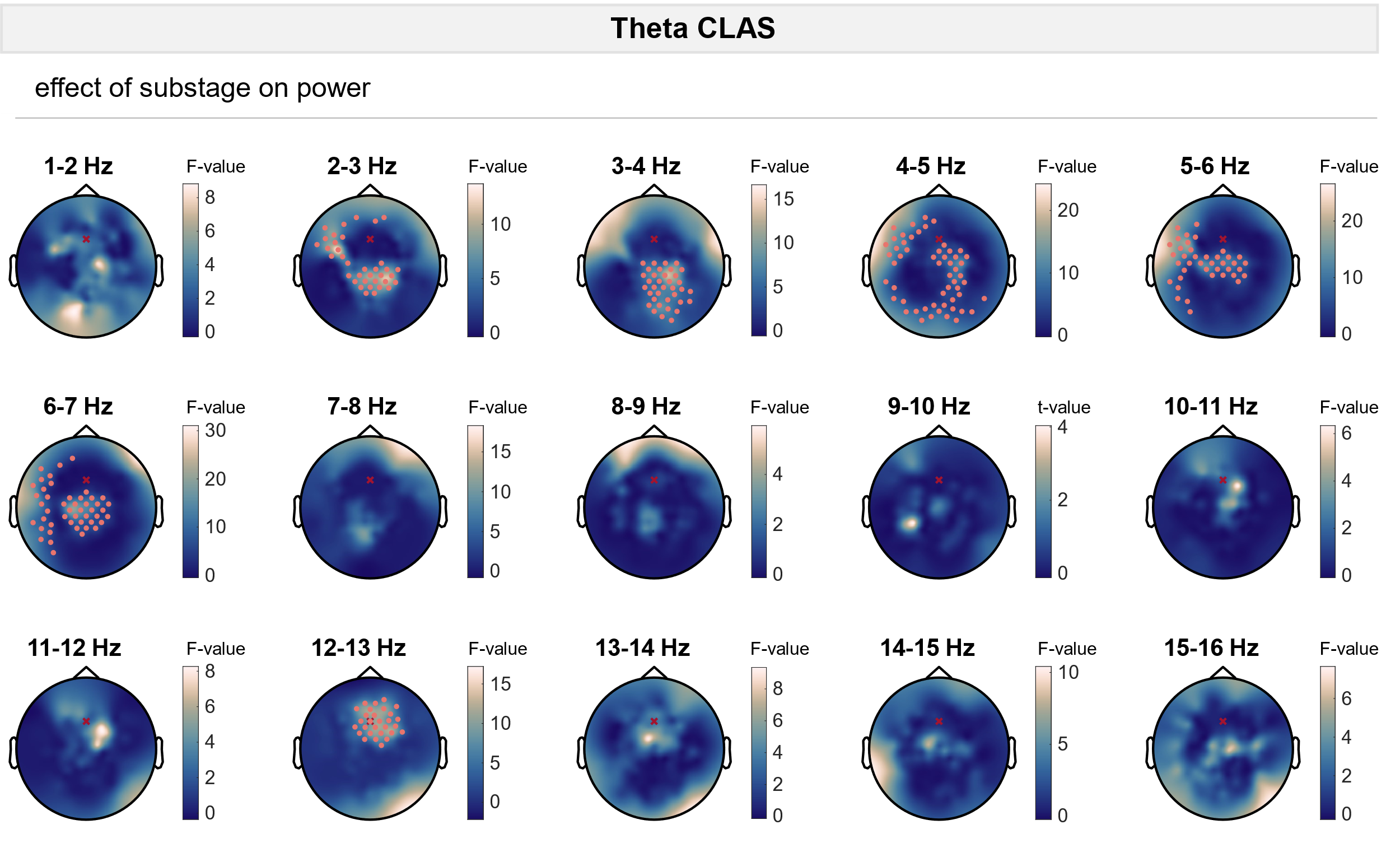

**Suppl. Figure 13.** To evaluate differences in power changes across phase conditions, for each 1-Hz frequency bin between 1 and 16 Hz and for each electrode, linear mixed-effects models with power change as dependent variable, phase condition, substage, and the interaction between substage and phase condition as fixed factors, and participant as random factor were calculated. F-values for the fixed factor substage are plotted for each electrode. Significant electrodes are indicated with pink dots (cluster-corrected). The electrode used for phase-locking is shown with a red cross.

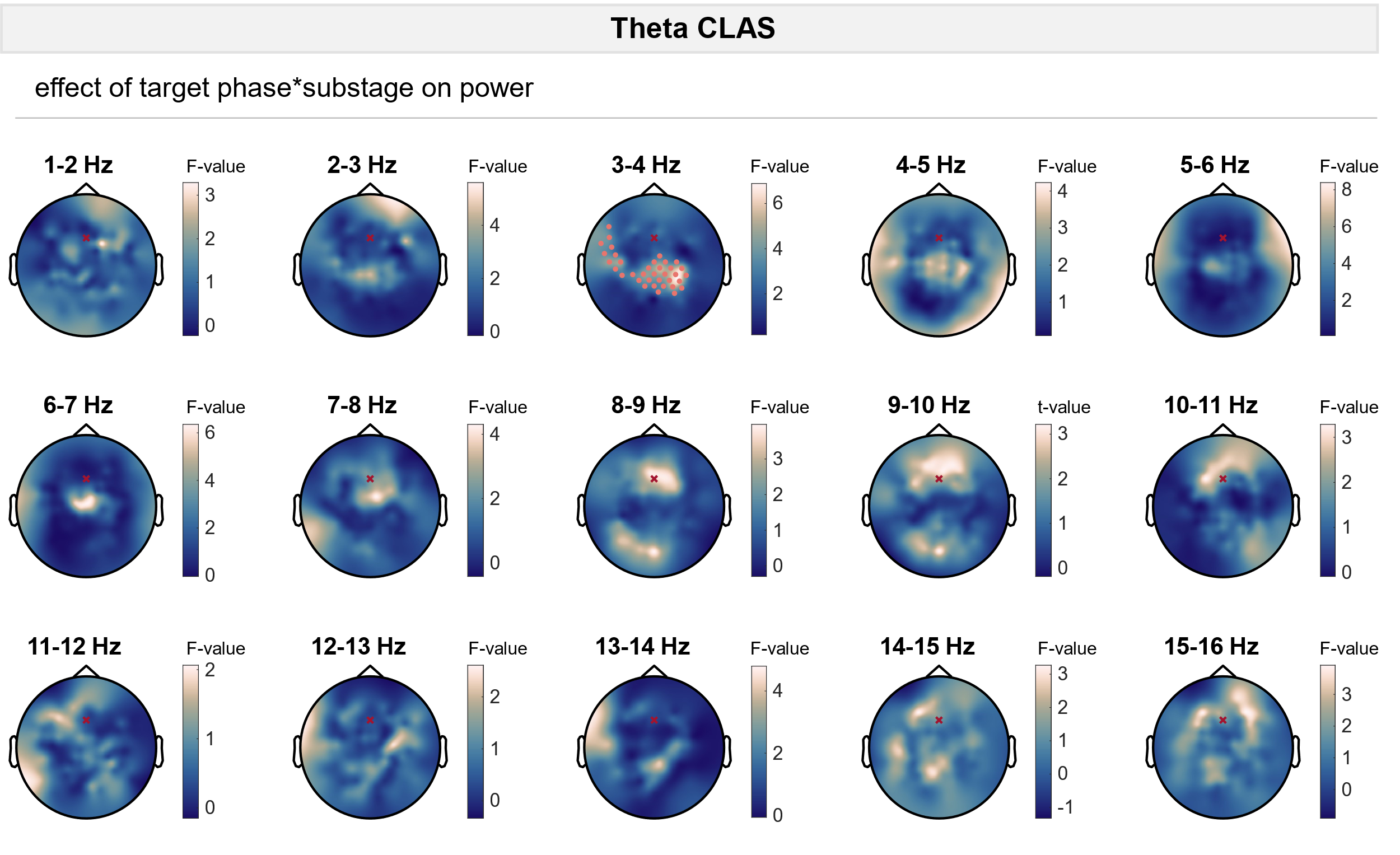

**Suppl. Figure 14.** To evaluate differences in power changes across phase conditions, for each 1-Hz frequency bin between 1 and 16 Hz and for each electrode, linear mixed-effects models with power change (ON vs OFF stimulation) as dependent variable, phase condition, substage, and the interaction between substage and phase condition as fixed factors, and participant as random factor were calculated. F-values for the interaction between phase condition and substage are plotted for each electrode. Significant electrodes are indicated with pink dots (cluster-corrected). The electrode used for phase-locking is shown with a red cross.

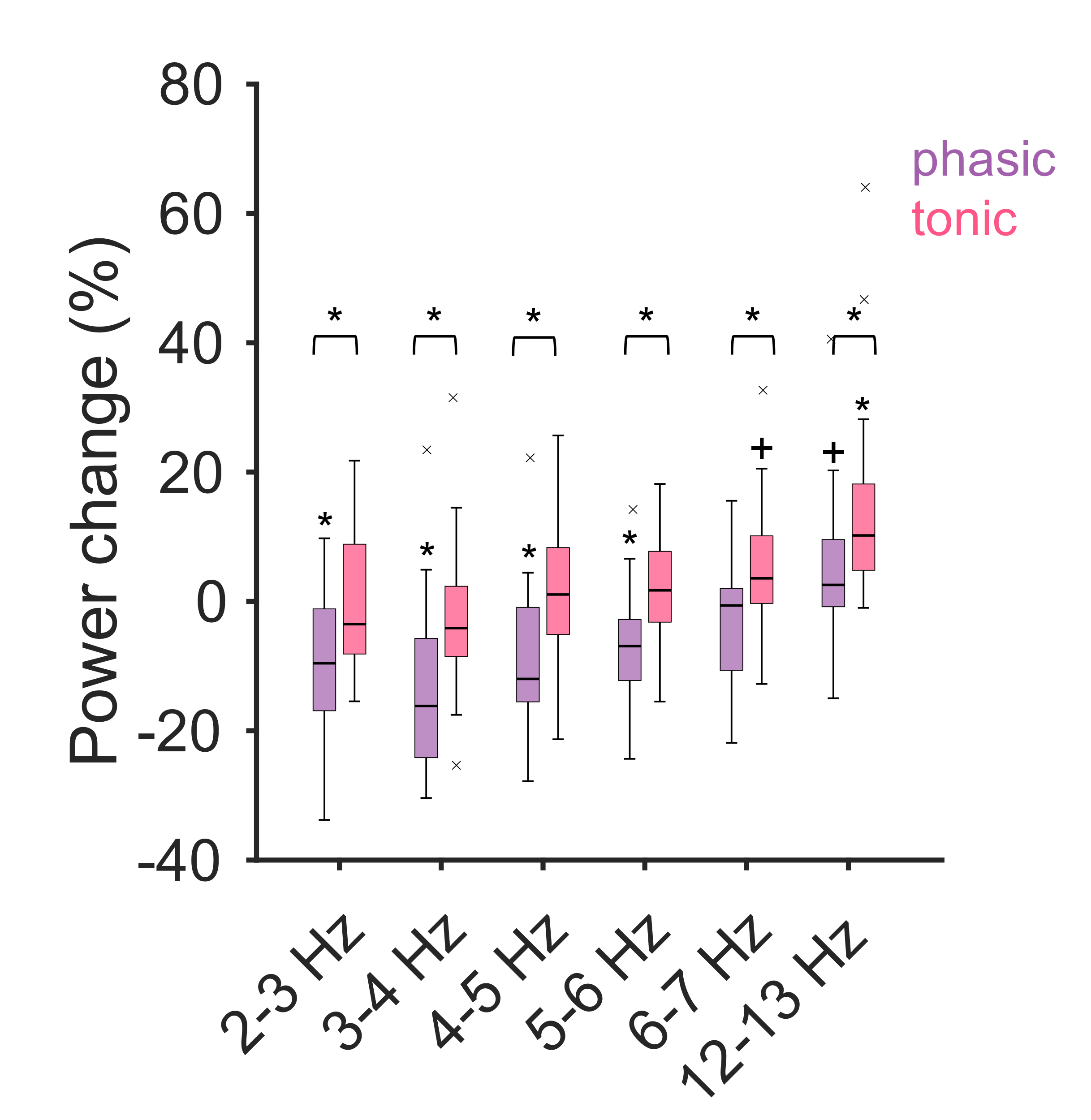

**Suppl. Figure 15.** Power changes for phasic and tonic REM sleep averaged across significant electrode clusters in **Suppl. Figure 13**. Significant differences between phasic and tonic REM sleep are indicated with black bars on top of the plot. Moreover, significant (p < 0.05) changes in a one-sample t-test are indicated with stars whereas trends (p < 0.1) are denoted with plus signs.

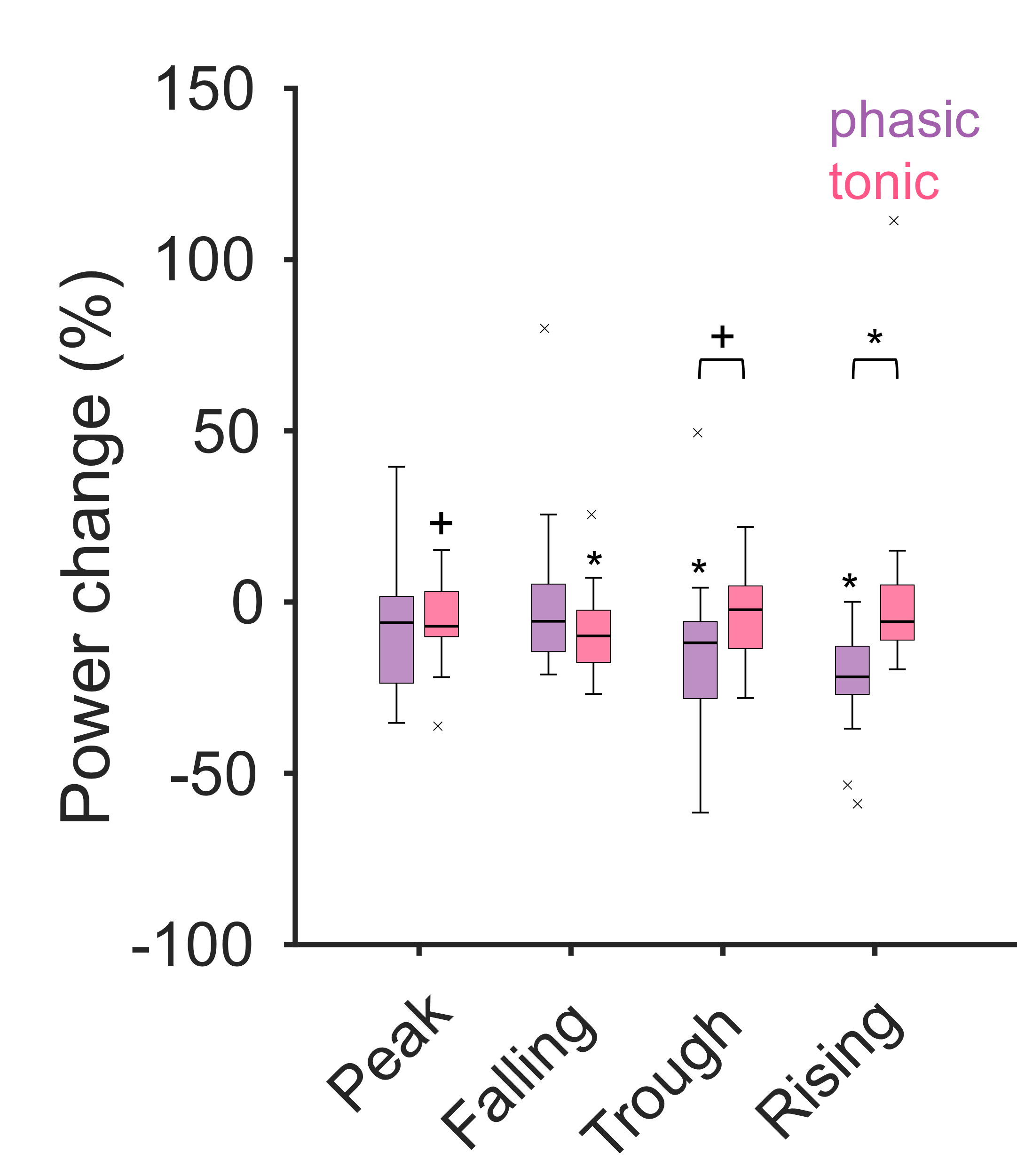

**Suppl. Figure 16.** Power changes for 3-4 Hz bin for phasic and tonic REM sleep and for different phase conditions averaged across significant electrode clusters in **Suppl. Figure 14**. Significant differences between phasic and tonic REM sleep are indicated with black bars on top of the plot. Moreover, significant (p < 0.05) changes in a one-sample t-test are indicated with stars whereas trends (p < 0.1) are denoted with plus signs.

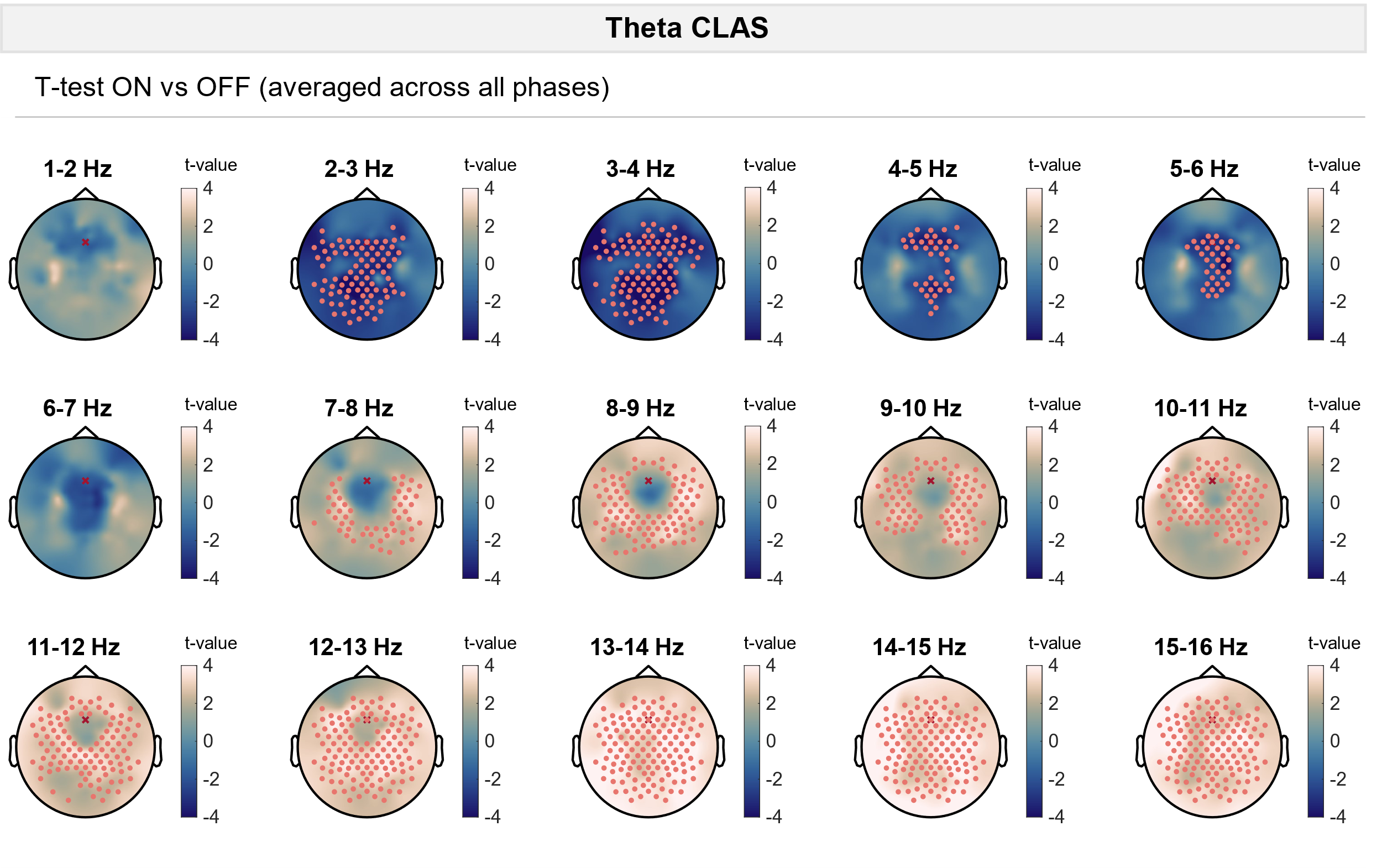

**Suppl. Figure 17.** To evaluate phase-independent effects of stimulation, power changes were averaged across all phase conditions for ON and OFF windows. For each electrode, power in ON and OFF windows was compared using a paired t-test. Significant electrodes are indicated with pink dots (cluster-corrected). The electrode used for phase-locking is shown with a red cross.

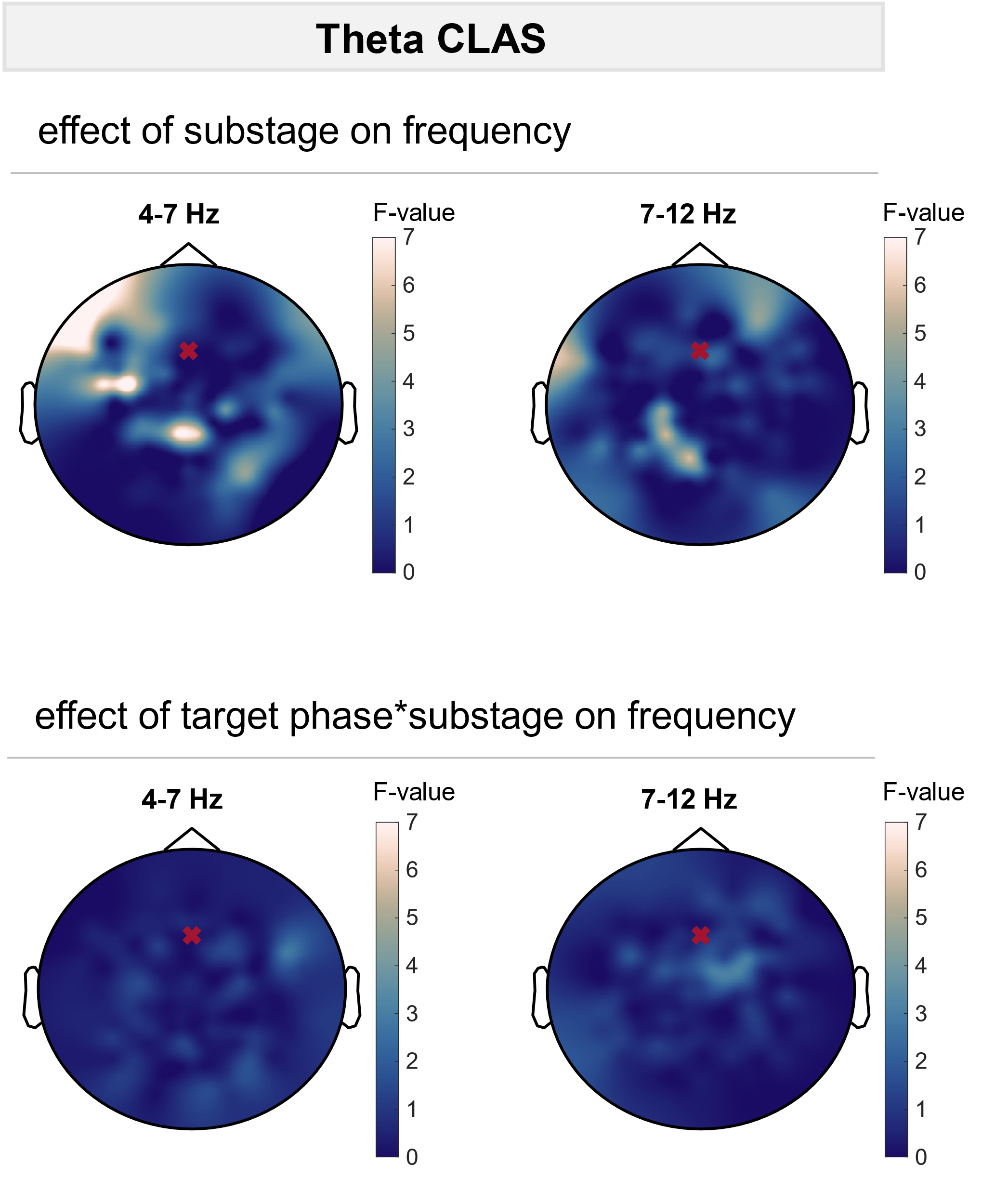

**Suppl. Figure 18.** To evaluate differences in frequency changes across phase conditions, linear mixed-effects models with frequency change as dependent variable, phase condition, substage, and the interaction between substage and phase condition as fixed factors, and participant as random factor were calculated. F-values for the fixed factor substage and the interaction between phase condition and substage are plotted for each electrode. No significant (p < 0.05) electrode clusters were found. The electrode used for phase-locking is shown with a red cross.

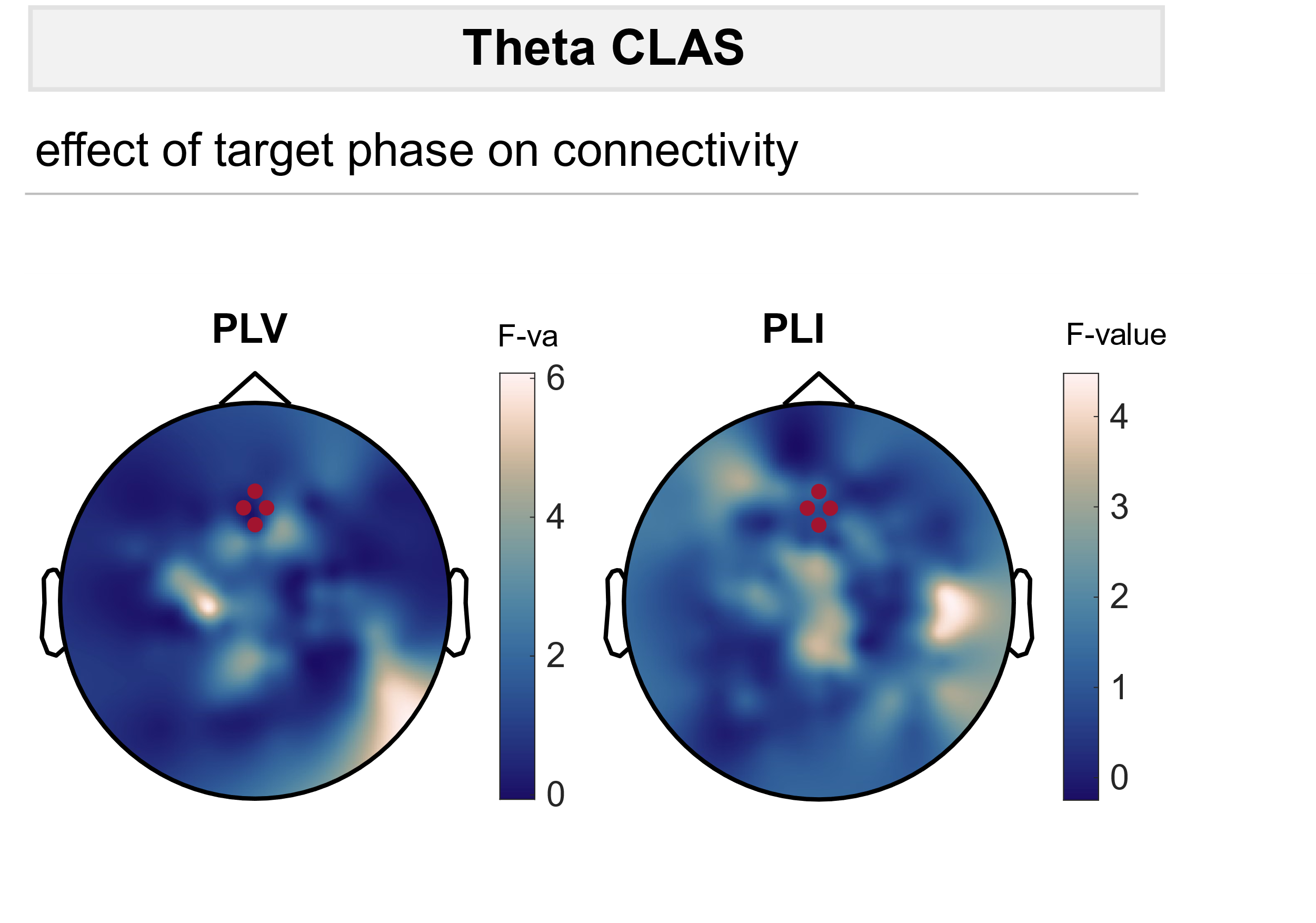

**Suppl. Figure 19.** Theta CLAS does not induce phase-dependent changes in connectivity. Connectivity changes were assessed by calculating the PLV between the targeted region i.e. the seed region (red dots) and all other electrodes. To evaluate differences in PLV and PLI changes across phase conditions, linear mixed-effects models with PLV or PLI change as dependent variable, phase condition, substage, and the interaction between substage and phase condition as fixed factors, and participant as random factor were calculated. F-values for the fixed factor phase condition are plotted for each electrode. No significant (p < 0.05) electrode clusters were found.
